# The condition of coral reefs in Timor-Leste before and after the 2016–2017 marine heatwave

**DOI:** 10.1101/2020.11.03.364323

**Authors:** Catherine JS Kim, Chris Roelfsema, Sophie Dove, Ove Hoegh-Guldberg

## Abstract

El Niño Southern Oscillation global coral bleaching events are increasing in frequency; however, the severity of bleaching is not geographically uniform. There were two major objectives of the present project: 1) assess the state of reefs and coral health at several sites and 2) explore water quality and climate change impacts on Timorese reefs. The impacts of climate change (principally by following coral mortality) were surveyed on coral reefs before and after the 2016–2017 underwater heatwave, using temperature loggers deployed between surveys which were compared to Coral Reef Watch (CRW) experimental virtual station sea surface temperature (SST). CRW is an important and widely used tool; however, we found the SST was significantly warmer (> 1°C) than *in situ* temperature during the austral summer accruing 5.79 degree heating weeks. *In situ* temperature showed no accumulation. Change in coral cover between surveys was attributed to reef heterogeneity. There were significant differences in coral cover, coral diversity, and nutrient concentrations between site and depth and a low prevalence of disease recorded in both years. The comparison of temperature and SST indicate that bleaching stress in Timor-Leste is potentially mitigated by seasonal and oceanographic dynamics. This is corroborated by Timor-Leste’s location within the Indonesian ThroughFlow. Timor-Leste is a climate refugium and the immediate conservation work lies in the management of localized anthropogenic impacts on coral reefs such as sedimentation and fishing.

## 1. Introduction

Timor-Leste is a developing country with limited infrastructure following decades of war and isolation. It is one of the poorest nations in East Asia representing one of six member-states of the Coral Triangle (CT), the global center of marine biodiversity (i.e., numbers of species), housing 29% of the world’s coral reefs [1,2]. Much of this diversity, however, is under threat due to a range of growing local and global stresses [3–6]. Globally, climate change-induced coral bleaching via ocean warming and coral disease are among the main threats facing coral reefs but are understudied in the CT compared to other reef regions. Furthermore, many coral diseases have been linked to increasing ocean temperatures, nutrient pollution, sedimentation, and fishing [7–10]. Global mass coral bleaching events or heatwaves, driven by anomalous increases in sea surface temperature (SST) maintained over time, have been occurring with increasing frequency [11]. Like disease, however, there is a paucity of data concerning the incidence and severity of bleaching in the CT. Additionally, these reefs are disproportionately threatened at a local level compared to other regions of the world [1].

### 1.1 Local threats to the coral reef of Timor-Leste

In addition to the threat posed by global climate change, there are a range of local impacts on Timorese reefs. Ninety two percent of reefs are at high or very high risk due to fishing pressure, watershed-based pollution, coastal development, and pollution from marine activities (i.e., shipping, oil and gas extraction) [1]. While the extent of destructive fishing practices has been decreasing since the Indonesian occupation from 1974 to 1999 [12], there are still an estimated 5,000 fishers that focus their fishing effort, without dynamite, on the narrow, productive shelf that supports coral reefs [13,14]. Fishing markets are limited to a very localized distribution given that the infrastructure for markets (e.g., refrigeration) is undeveloped [15]. Additionally, gleaning, or harvesting invertebrates from intertidal flats for consumption, known locally as *meti*, is commonly practiced by women and children and has its own, often significant, impacts [15,16]. Similarly, agricultural practices are generally limited to small-scale, subsistence farming without the use of non-organic fertilizers and pesticides although the development of such practices is outlined to improve food security [17].

Watershed-based pollution is widespread due to deforested landscapes that lead to large volumes of unsettled sediment and pollution flowing downstream and into coastal waters. An estimated 24% of forests in-country have been lost from 1972 to 1999 due mostly to slash and burn agriculture and logging during the Indonesian occupation and because of its importance as fuel [18–20]. Significant development is planned over the coming decades with potential increased coastal impacts [1,18–20].

### 1.2 Disease in the context of coral reef health

While rapid ocean warming has increased mass coral bleaching and mortality [21], other consequences of stress have been increasing including the prevalence of coral disease. Coral disease has been a major contributor to the decline of corals in other regions such as the Caribbean [22], and also severely threatens reefs in the Indo-Pacific [4,23–25]. By contrast, there have been relatively fewer studies of coral disease in the CT (Table A1) [26]. In this study, diseases were defined as syndromes caused by pathogens and recorded abiotic diseases such as coral bleaching under the broad category of compromised health [26].

Disease and other signs of compromised physiology are one of many indicators of condition of coral reefs (loosely defined as coral health). Understanding the signs of declining coral condition has the potential to alert reef managers to potential problems (i.e., a change in the level of local threats). Therefore, it is important to document lesions, or morphologic abnormalities, predation, physical breakages (i.e., storms, anchors), and aggressive interactions which may result in tears or breaks in the tissue, partial mortality, and stress to the coral host. Disease can be endemic and highly visible [22], or present in low frequency in any given population [25]. Tracking disease and other signs of compromised health through time can be paired with other datasets (i.e., herbivore biomass, hurricane incidence, environmental parameters, etc.), and is related to key physiological parameters such as growth rates, fecundity, and community composition of reefs [27]. At most sites in Timor-Leste, these types of measurements are absent, highlighting the importance of the present study as a crucial baseline on the conditions of important marine resources.

### 1.3 Water quality and coral reefs long the north coast of Timor-Leste

Pollution arising from disturbed coastal regions and watersheds poses a serious threat to coral reefs. This type of pollution includes a wide range of compounds such as agrichemicals (i.e., pesticides), inorganic nutrients (i.e., nitrate, ammonia, and phosphate), soils and sediments, and fossil fuel residues that flow from disturbed landscapes. Many of these compounds negatively affect coral physiology by reducing calcification rates, fecundity, fertilization success, and larval development [28]. This can degrade reef communities, reducing coral cover, community composition diversity, and structural complexity [29,30]. High levels of marine pollution can increase the prevalence and severity of disease and susceptibility to bleaching [31–35]. Dissolved inorganic nitrogen (DIN = ammonium + nitrate + nitrite) measurements on reefs are generally < 1.5 μM (individual species ammonium, nitrate, nitrite < 1 μM) with lower phosphate concentrations (< 0.3 μM; Table A2) [36–43]. A greater prevalence of disease has been associated with elevated concentrations of DIN from anthropogenic sources (i.e., fertilizer, sewage pollution, etc.) and phosphate ranging from 3.6 μM to 25.6 μM and 0.3 μM to 0.4 μM respectively [36,37,41–43].

The isotopic signature of nutrients such as nitrogen can often act as a tracer for different sources of coastal pollution with different forms having different impacts (e.g., sewage can increase pathogen concentrations) and solutions [10,44–53]. Stable isotope analyses of nitrogen stored in macroalgae can provide a nutrient signal integrated over time versus water sampling, which is highly variable over space and time [54]. Generally, δ^15^N signatures in algae associated with urban wastewater are > 10‰ [55–58]; however, values as low as 4.5‰ have been argued to be a result of anthropogenic sources of nutrients [47]. Depleted δ^15^N values (1–3.5‰) can be sourced from either synthetic fertilizers [55,57] or pristine mangroves [59]. Natural and synthetic fertilizers display a large range from −4‰–+4‰ of δ^15^N values while nitrogen fixation typically has a negative δ^15^N signature between −2–0‰ [60]. Upwelling can have variable δ^15^N values ranging from 5–12‰ [46,59,61–64]. Given the lack of inorganic fertilizer use and waste infrastructure in Timor-Leste, nearshore waters were expected to have δ^15^N signatures higher than upwelling (5-6‰) which is indicative of sewage pollution (> 10‰). Both fertilizer use and waste infrastructure are expected to be developed as described in the national strategic development plan [17].

### 1.4 Global Impacts – ocean warming, mass coral bleaching, and mortality

The mass global bleaching event in 2016–2017 was the longest and most intense in history [21,65]. This El Niño Southern Oscillation (ENSO) associated thermal event had global, but patchy impacts on coral reefs. Few reports exist of the impacts in the CT. The CT arguably, however, has the most to lose from the degradation of reefs [1]. NOAA’s Coral Reef Watch virtual station in Timor-Leste (CRWTL) reported anomalous warming between the two survey periods of November 2015 and July 2017. Between January and May in 2016, and again from January and February 2017, the water temperature of the regions attained degree heating weeks (DHWs) above 4, but less than 8 [66]. A DHW range of 4 to 8 has been associated with 30–40% bleaching [67,68], suggesting that corals may have bleached twice within the 20-month sampling interval. Surviving corals, however, would have had four to five months to recover before resurveying in July 2017. Typically, mortality is not expected below DHW of 8 [69], although this is variable between species [70,71]. DHWs of or above 8 were not attained in Timor-Leste during the experimental period. Corals that have experienced a recent thermal event that is sufficiently warm to cause temporary bleaching in some corals, may nonetheless be vulnerable to disease or other signs of compromised health [4,72,73]. Additionally, corals may endure sublethal effects for months after the event as they attempt to rebuild energy reserves [5,74]. During the 2017 bleaching event on the Great Barrier Reef, 48% of tabulate Acroporids were co-infected with White Syndrome (WS) and had seven times more tissue loss than only bleached colonies [75].

The aims of the present study were two-fold. The first was to investigate the state and health of coral reefs as measured by the presence of coral disease and other signs of compromised health. The second aim was to explore the impacts of humans on Timorese reefs through water quality measurements and surveys before and after the 2016-2017 global bleaching event. This was achieved through repeated coral health surveys, seawater nutrient and nitrogen stable isotope analyses of macroalgae to assess nutrients, and *in situ* and remotely sensed temperature data.

## 2. Materials and Methods

### 1.1 Study Site

Timor-Leste is a small country inside the southern edge of the CT and between Australia and Indonesia. The country gained its independence in 2002 following nearly 25 years of Indonesian occupation. It lies within the Indonesian ThroughFlow (ITF), a major oceanographic feature connecting the Pacific and Indian Oceans [76]

This study was undertaken along the coast of Dili, Timor-Leste to complement a growing body of coral reef science in the area. Previous indications of reef health in this area have typically been anecdotes from surveys with other objectives. Dili, the capital (8°33’S and 125°34’E), houses a quarter of the country’s population with 252,884 people recorded in the 2015 Census [77]. The seasonal Comoro River runs through Dili, with flow ranging from less than 0.5 m^3^/s from July to November to 12.3 m^3^/s in March during the monsoon season from December to May [18,78]. The present study was conducted in two, three-week field trips that occurred in November of 2015 and July of 2017 during the dry season. The dry season offers safer surveying conditions but would also limit terrestrial run-off inputs such as nutrients. While future studies should expand the results here by examining the dynamics of coastal systems during the wet season, it was not investigated here.

Surveys were conducted at four sites. Two sites flanked Dili and were representative of reefs under urban influences (“Urban-W” with 5,017.9 people/km^2^; “Urban-E” with 779.5 people/km^2^) and two sites were representative of reefs under rural influences (“Rural-N”, and “Rural-E”; Figure 1). Sites were chosen for logistics and to complement US National Oceanic and Atmospheric Administration climate station data collection surveyed between 15–27^th^ of November 2015 and 15– 29th of July 2017 [79].

**Figure 1.**
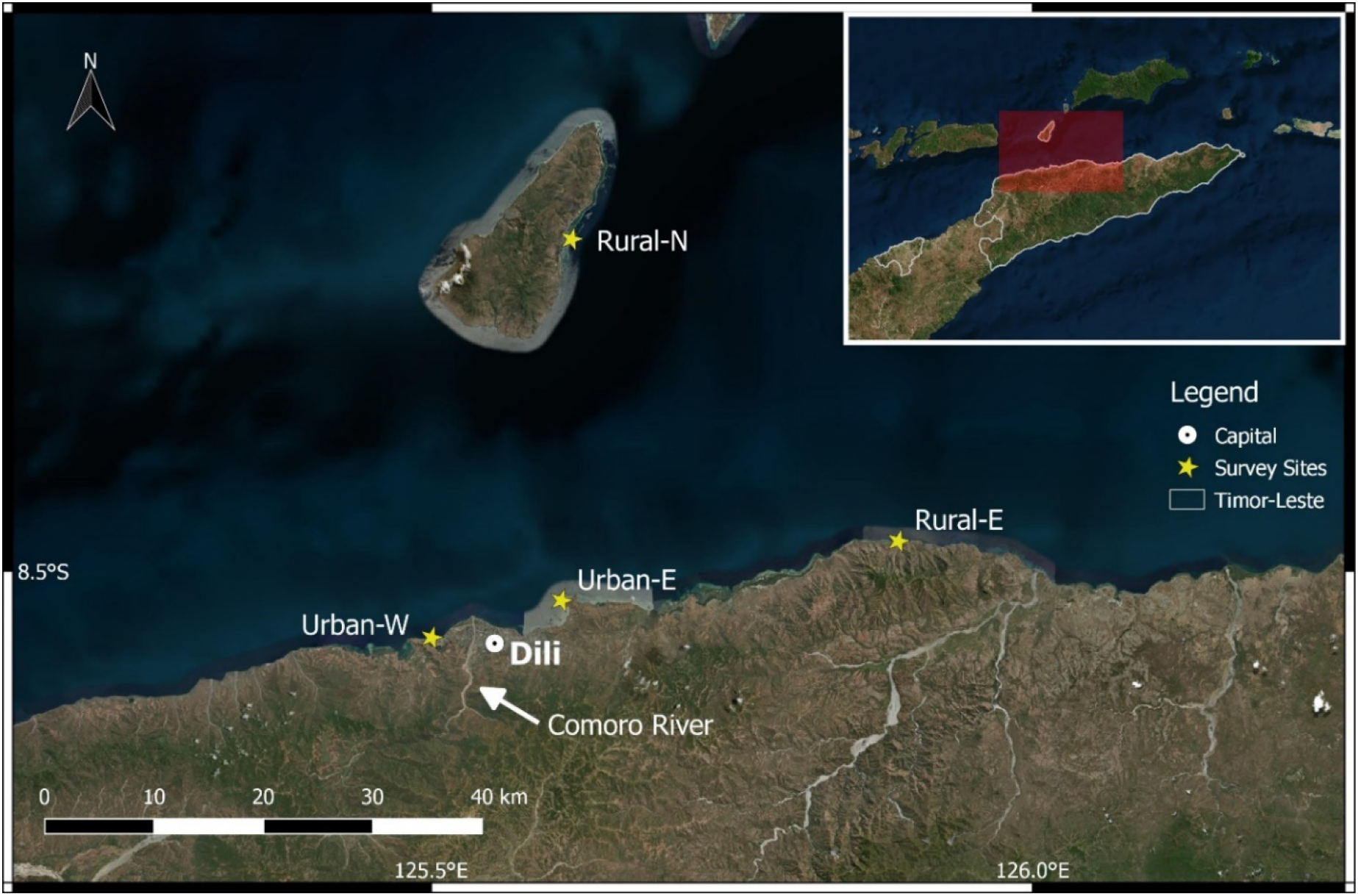
Survey sites in Timor-Leste around the capital of Dili. Rural-N on Ataúro Island in the channel, Rural-E 40 km east of Dili, and Urban-W and Urban-E flanking Dili. The highly seasonal Comoro river can be seen just east of Urban-W. The four sites were sampled at two-time points in November 2015 and June 2017. Jaco Island lies on the easternmost point of the country.

### 2.1 Coral community composition and coral health surveys

To assess benthic cover and coral health we deployed 15 m line intercept transects [80] and 15 m × 2 m belt transects [27]. At each of the four sites, three transects were laid at 5 m (reef flat) and 10 m (reef slope) depths for a total of 24 transects across all sites. The first transect at each site was chosen randomly with the subsequent transects at least 5 m away at the appropriate depth contour. For the line intercept transect, the benthos under the 15 m tape was categorized into a major benthic category (i.e., hard coral, soft coral, substrate/sand, macroalgae, turf algae, cyanobacteria, and crustose coralline algae-CCA). On the coral health belt transects, every coral colony within the belt transect area was identified to genus and assessed visually for coral disease and signs of potential compromised coral health consisting of overgrowth by macroalgae, turf and cyanobacteria, encrusting invertebrates (sponges, tunicates, flatworm infestation), burrowing invertebrates (worms, barnacles), signs of predation (fish and *Drupella sp.* snails), signs of bleaching (partial or total loss of algal symbionts appearing white), signs of coral response (pigmentation, mucus), and physical damage (sedimentation, breakage) as per protocols developed by the Global Environment Facility and World Bank Coral Disease Working Group (Figure 2; Figure A1; Table A3) [27]. Any uncertain diagnoses were photographed and used in later consultation. The prevalence of disease and compromised health was calculated as the number of corals affected by disease/compromised category divided by the total number of corals in the transect. The same transect start GPS coordinates at the surface were used for the second survey, in July 2017, with the same direction considering currents, etc.

**Figure 2.**
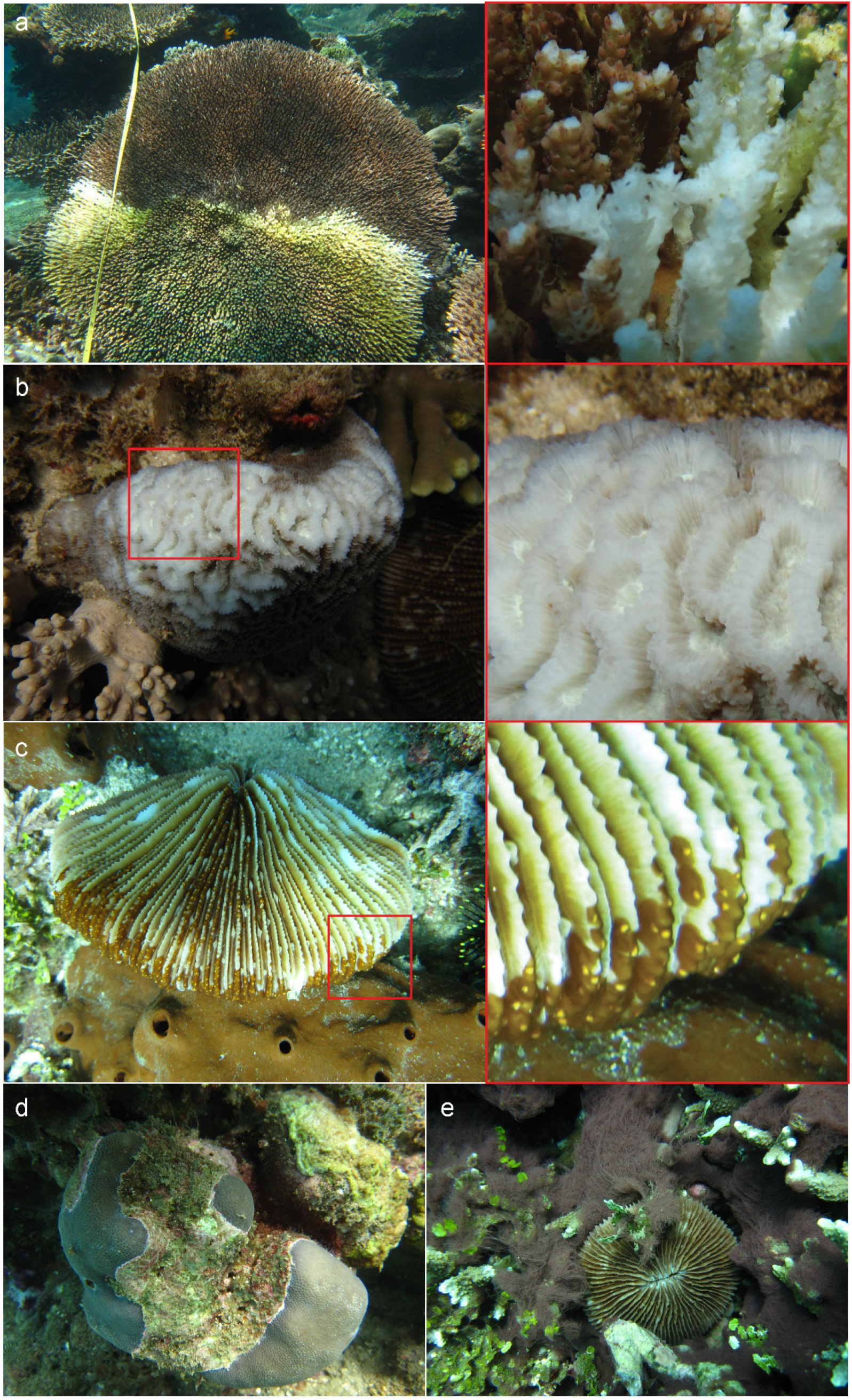
Examples of disease and compromised health observed during the surveys undertaken in Timor-Leste between November 15-27^th^, 2015. a) WS–White Syndrome band of distinct tissue loss on tabulate Acroporids with white skeleton abutting live tissue with exposed skeleton gradually colonized by turf algae, b) bleaching, c) flatworm infestation, and d) turf algae overgrowth. See Figure A1 for other compromised states and Table A3 for more information.

### 2.2 Measurement of nutrient concentrations and stable isotope ratios

Seawater samples were collected for measuring the concentration of inorganic nutrients as an indicator of nutrient pollution. Three replicate 100 ml seawater samples were collected on each transect 0.5 m above the benthos and kept on ice until filtering through a 0.22 μm pore membrane filter and stored frozen. Seawater samples were analyzed within four months for ammonium, nitrite, nitrate, and phosphate using flow injection analysis at the Advanced Water Management Center (The University of Queensland). Nitrite had mostly zero values and was combined with nitrate for analyses.

Macroalgal samples were collected for stable isotope analysis to explore the origin of inorganic nitrogen. Three replicates of *Halimeda sp.* and *Chlorodesmis sp.* macroalgae (approximately 5 g dry weight) were collected when found on each transect, rinsed, and air-dried for transport. In the laboratory, the samples were re-dried at 60°C for a minimum of 24 h before homogenization using a mortar and pestle and analysis at the Cornell University Stable Isotope Laboratory (Finnigan MAT Delta Plus isotope ratio mass spectrometer) for δ^15^N analysis.

### 2.3 In situ and satellite temperature data

HOBO pendant temperature loggers (Onset Computer Corporation, Bourne, MA USA) were deployed at every site and depth in November 2015 recording temperature every 30 min. All were retrieved except those from Rural-E in June 2017. Remotely sensed satellite SST data from the NOAA’s CRWTL was downloaded from August 2015 through August 2017. This product uses 5 km^2^ resolution to predict bleaching stress across an entire jurisdiction such as Timor-Leste versus values at every pixel [66].

### 2.4 Statistical analyses

All analyses were conducted in R version 3.6.3 [81] and PRIMER7 [82,83]. Repeated measures permutational multivariate analysis of variance (PERMANOVA) with 9,999 permutations were conducted to test for significant effects between sites (Rural-N, Rural-E, Urban-W, Urban-E), depths (5 m, 10 m), and years (2015, 2017) on a Bray-Curtis similarity matrix of transformed benthic cover categories, transformed prevalence of disease and compromised health, and zero-adjusted, transformed Bray-Curtis similarity matrix of the number of colonies per coral genera (i.e., the count of coral genera per transect) [82,83]. All multivariate tests were also tested for homogeneity of dispersion akin to the homogeneity of variance in univariate tests [82]. Repeated measures analysis of variance (ANOVA in the *car, emmeans, nlme* R packages) [84–86] was used to test transformed hard coral, categories of disease and compromised health (bleaching–square root transformed), the transformed number of coral genera, and Shannon diversity index on transformed coral genera for significant effects between sites, depths, and years. Principal coordinates analysis (PCO) was run on the same transformed resemblance matrix of coral genera to visualize coral community structure. A repeated measures ANOVA was also conducted on the log-transformed number of Acroporids per transect between site, depth, and year. Normality was visually inspected (*hist, qqplot, qqnorm*, *leveneTest* in the *car* package) and all previous transformations were square root.

Nutrient data were only collected in 2015 and a two-way ANOVA with factors, site, and depth was performed on the seawater nutrient data including DIN (transformations: log–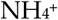 and DIN, square root-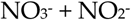). A two-way ANOVAs (*Anova*) was used to test for significant differences in δ^15^N for each of the two genera of algae, *Halimeda sp*. and *Chlorodesmis sp*., with the factors site and depth. Only three samples of *Chlorodesmis sp.* were collected on a singular transect at Rural-E and were removed from the analysis. Variables were visually inspected for normality and tested for homogeneity of variance using Levene’s test (*leveneTest*). Percent nitrogen was log-transformed for *Halimeda sp*. Posthoc tests were conducted (*multcomp* and *emmeans* R packages) for *Halimeda sp*. and *Chlorodesmis sp*. respectively [87]. The effect of 2015 nutrients on the same resemblance matrix of the prevalence of disease and compromised health of the same year was analyzed using a PERMANOVA (9,999 permutations) with site and depth as factors and covariates 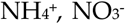, and 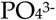 from the collected seawater data.

The monthly average from the moving seven-day average of the 24 hr daily maximum temperature of *in situ* temperature logger data and remotely sensed CRWTL data over the same time was calculated. ANOVAs were used to test for differences in site, depth, and month nested in year between monthly means of in situ temperature logger data and season and site between the average of CRWTL temperature over the study period and all logger sites over the following austral seasonal groupings: Jan–Mar (summer), Apr–Jun (fall), Jul–Sept (winter), Oct–Dec (spring). To assess levels of thermal stress, *in situ*, DHWs retrieved from CRWTL online [66].

## 3. Results

Benthic community composition and coral cover varied significantly between site, depth, and year and there was a trend of rural sites having more live coral; however, a greater number of sites are needed to draw this conclusion. The changes in benthic composition between surveys were attributed to heterogeneity versus coral mortality following the 2016–2017 global bleaching event. This was supported by the *in situ* temperature data collected between surveys which never surpassed maximum monthly mean (MMM) + 1°C to accumulate DHWs. Conversely, the CRWTL SST product accumulated 5.79 DHWs over the same time. The underlying coral community structure was significantly different at the four sites and significant differences in variability between and within sites contributed to this effect which can be a sign of varying levels of impact. There was a low prevalence of disease at the four sites surveyed in Timor-Leste which were not related to whether sites were urban versus rural. Contrary to our hypothesis, WS was the most prominent at Rural-N with a prevalence of 0.9 ± 0.2% in 2015 while GAs was the most prevalent at Urban-W in 2017 (0.6 ± 0.3%). Lastly, seawater nutrients and δ^15^N were not significantly elevated at urban sites or consistently greater at 5 m depth versus 10 m. The prevalence of disease was significantly associated with phosphate concentrations as the highest combined nitrate and nitrite and phosphate were documented at Rural-N at 10 m, the site of highest disease prevalence.

### 3.1 Coral cover and community composition at four sites

A total of 9,521 corals of 51 genera were counted over 720 m^2^ of the surveyed area per year in 2015 and 2017. The benthic composition was significantly different between the four sites. Rural sites had higher coral cover than the urban sites and the overall patterns of coral cover were consistent across survey years (Figure 3). Coral diversity also varied significantly; however, lower or higher genera diversity did not fall along rural versus urban distinctions. Urban-W at 10 m had low coral diversity while Rural-N was the only site dominated by tabulate and branching Acroporids and consistently high (> 40%) coral cover over survey years.

**Figure 3.**
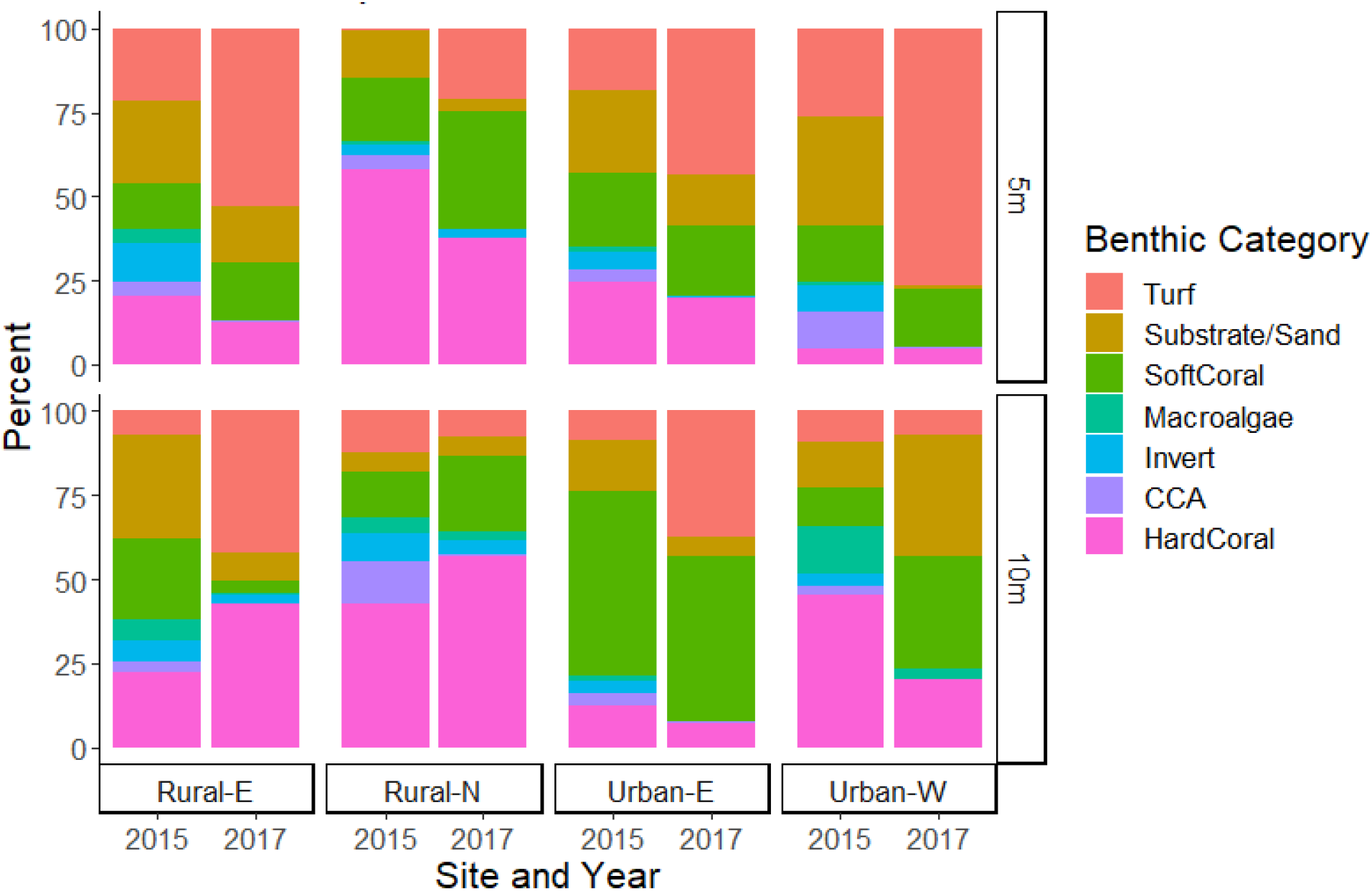
Benthic composition cover from 15 × 2 m line intercept transects by site and depth for 2015 and 2017 survey periods in Timor-Leste. Major categories include Hard Coral, CCA–crustose coralline algae, Invert-mobile invertebrates, Macroalgae, Soft Coral, Substrate/Sand, Turf Algae.

Benthic composition varied spatially with a significant site and depth interaction (repeate measures PERMANOVA, pseudo-F(3,47) = 4.5117, p(perm) = 0.0041) and temporally by year (pseudo-F(1,47) = 34.0270, p(perm) = 0.0002). At 5 m depth, Rural-E had comparable benthic composition to both urban sites but was only similar to Urban-W at 10 m (p > 0.05). Urban-W was the only site that varied significantly between depths (p < 0.05). Coral cover was significantly different with a three-way interaction (repeated measures ANOVA χ^2^ (3,16) = 13.6947, p = 0.0034). Urban-W at 5 m had the lowest coral cover in both years (mean ± SE; 4.8 ± 1.8% in 2015 and 4.5 ± 1.5% in 2017) and Rural-N 5 m (58.2 ± 1.7%) and Rural-N 10 m (56.9 ± 3.3%) had the highest live coral cover respectively in 2015 and 2017 (Figure 5). Overall, hard coral cover was higher at rural sites (37.3 ± 5.3%) than urban sites (12.9 ± 3.8%).

Although 51 genera were found across the four sites, few genera dominated the reef, namely *Porites* (2015 = 17.4%, 2017 = 13.0%, *Fungia* (2015 = 13.7%, 201 = 19.0%), and *Montipora* (2015 = 12.9%, 2017 = 13.4%). The maximum genera richness of 33 ± 2 was present at Rural-N and minimum at Urban-W at 10 m with 18 ± 2 genera. The Shannon diversity index showed site and depth differences (three-way repeated measures ANOVA χ^2^(3,16) = 24.3377, p < 0.0001) with Urban-W 10 m (1.7 ± 0.2) driving this interaction (Figure A2). The coral diversity was similar across rural and urban sites except for Urban-W at 10 m.

Coral diversity, as measured by the abundance of individual coral genera, also differed significantly by a site and depth interaction which was driven by site-level distinctions versus rural and urban boundaries (repeated measures PERMANOVA pseudo-F(3,47) = 3.3011, p(perm) = 0.0018). Diversity at Urban-E was significantly different from all other sites (p < 0.05) at 5 m and all sites were significantly different at 10 m (p < 0.05). Rural-E and Urban-W were significantly different within sites between depths (p < 0.05). Sites were generally distributed along axis two of the PCO with Rural-N most positively associated with tabulate Acroporids, *Galaxea*, *Goniastrea*, *Montipora*, and *Stylophora* genera, while 5 m transects were aligned along axis one with more *Pocillopora*, *Platygyra*, and massive *Porites* corals (Figure 4). Dispersion, or variability within the coral genera, was also significantly different for the site and depth interaction (F = 10.638, p(perm) = 0.0001) indicating that variability within sites and depths contributed to significant differences in addition to abundances of coral genera. Specifically, the dispersion was significantly lower at Rural-N compared to Urban-E and Urban-W at 10 m and greater at Urban-W 10 m compared to the same site at 5 m, Urban-E 10 m, and Rural-E 10 m (p < 0.05). Site dispersion, or spread of site markers, increases moving down PCO axis two and is generally less at 5 m (open symbols) than at 10 m (solid symbols; Figure 4).

**Figure 4.**
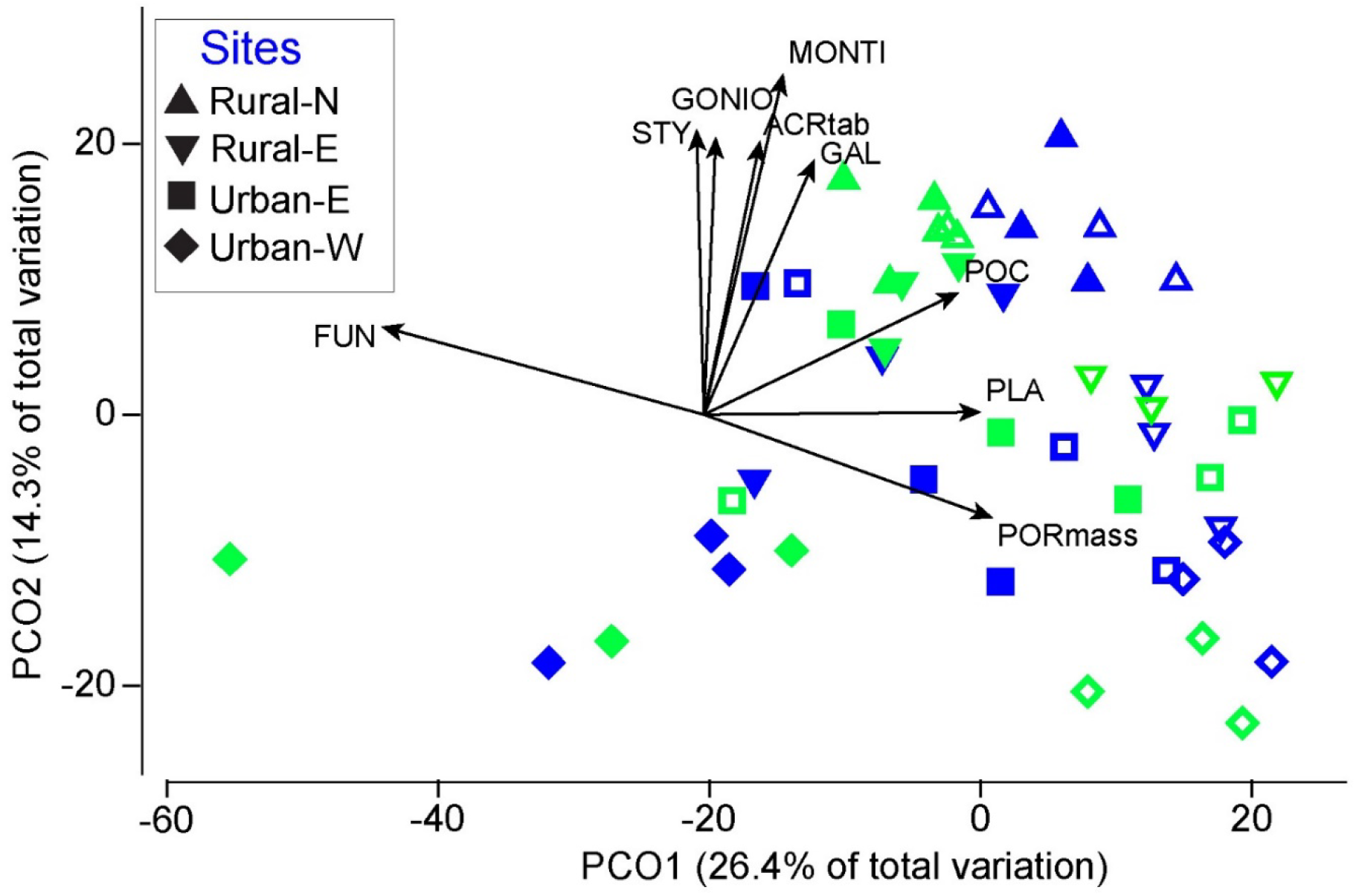
Principal Coordinates Analysis biplot of coral genera diversity. Shapes indicate site with empty and solid markers indicating 5 and 10 m depths respectively. Color indicates survey year: blue– 2015 and green–2017. Abbreviations are coral abundances as follows: ACRtab–Acropora tabulate, FUN–Fungiids, GAL–Galaxea, MONTI–Montipora, GONIO–Goniastrea, PLA–Platygyra, POC– Pocillopora, PORmass–Porites massive, and STY–Stylophora.

**Figure 5.**
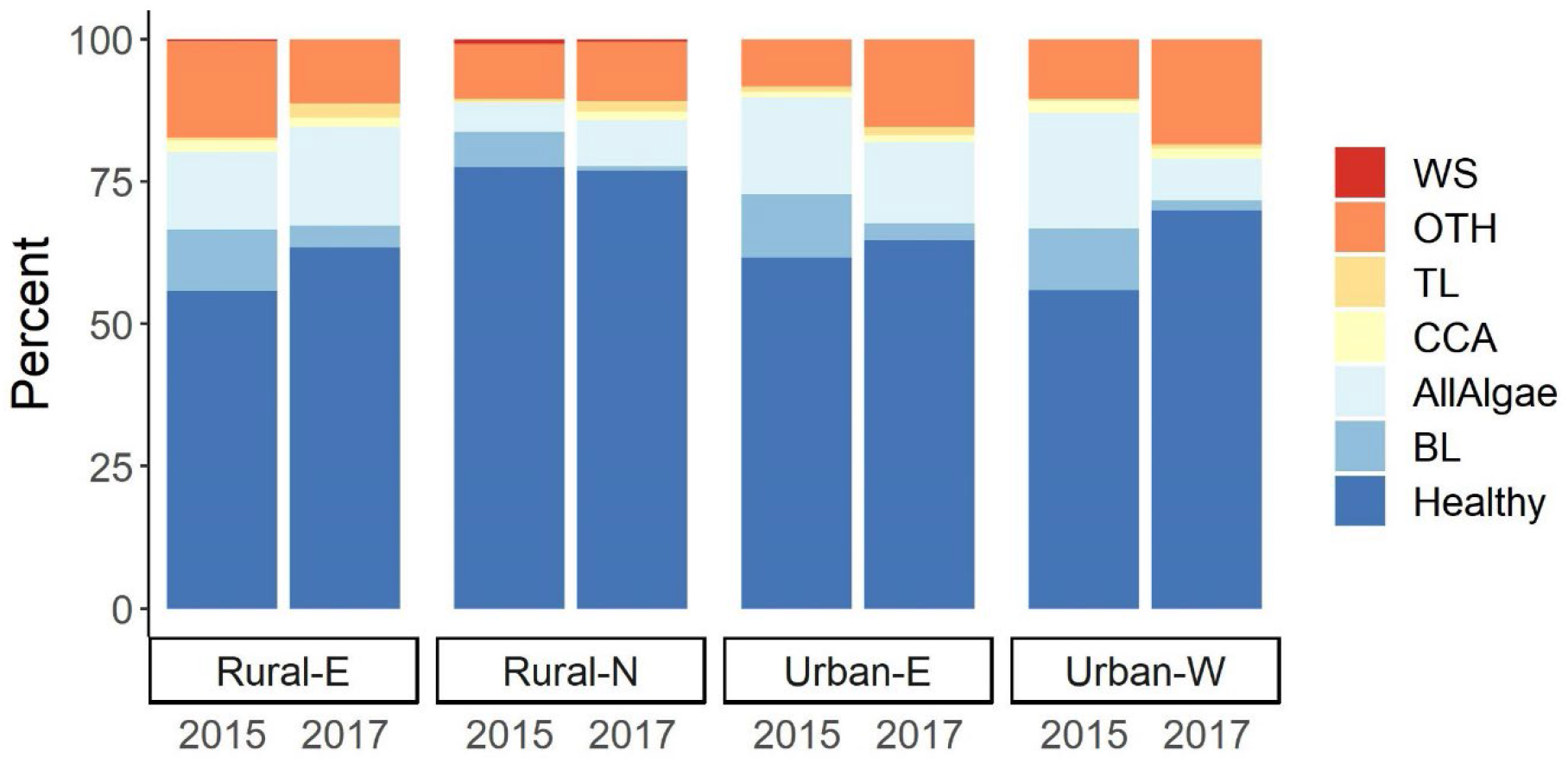
Prevalence of disease and indicator of compromised coral health from 15 × 2 m belt transect surveys at four sites in Timor-Leste from November 15-27^th^, 2015 and June 15-29^th^, 2017. AllAlgae– combined macroalgae, turf, and cyanobacteria overgrowth; BL–Bleaching; CCA–Crustose coralline algae overgrowth; OTH–combined pigmentation, predation, invertebrate infestation/overgrowth, burrowing invertebrates; TL–Unexplained tissue loss; and WS–White Syndrome.

There was also a significant site and year interaction for coral community structure (repeated measures PERMANOVA pseudo-F(3,47) = 2.1432, p(perm) = 0.0071). Coral diversity between Rural-N and all other sites and Urban-W and Rural-E were significantly different in both years (p < 0.05) and differences in dispersion were also significant where Rural-N and Urban-W had the least and most variability respectively. Dispersion at these sites was significantly different from all other sites for both in years in the case of Rural-N and 2015 for Urban-W (Figure 4).

Community composition varied significantly between sites with sites invariably having different dominant genera. While rural sites had more coral cover overall, coral community composition was distinct between sites at differing levels between depths. Urban sites did have low coral cover (< 10%) at either 5 or 10 m consistently between years (Figure 3). Additionally, Urban-W had the lowest coral diversity at 10 m (Figure A2). Rural-N stood out with the highest coral cover, the greatest number of genera, and the largest proportion of tabulate and branching Acroporids. At both depths, Rural-N had significantly more tabulate (repeated measures ANOVA χ^2^(3, 20) = 88.7746, p < 0.0001) and branching Acroporid colonies (repeated measures ANOVA χ^2^(3, 20) = 38.3591, p < 0.0001) than all other sites with 21.1 ± 0.7 and 11.0 ± 0.4 colonies per transect respectively (p < 0.05). All other sites averaged less than five Acroporid colonies per transect for both morphologies.

### 3.2 Prevalence of coral disease and indicators of compromised health

Overall, the majority of hard corals at sites surveyed appeared healthy. Those categorized as “healthy” made up 65.7 ± 1.70% of corals surveyed averaged over both years with low (< 1%) prevalence of diseases. There were no clear distinctions between rural and urban sites. In 2015, there was 0.9 ± 0.2% prevalence of WS at Rural-N with 61.9% of total disease found on *Acropora sp.* Rural-N also had the highest prevalence of GAs the same year with 0.6 ± 0.2%. There was one case of unconfirmed Trematodiasis, which requires microscopic confirmation of the larval trematode. Disease prevalence was lower in 2017 with the highest prevalence of WS at Rural-N again (0.5 ± 0.1%) but, Urban-W had the most GAs (0.6 ± 0.3%). All cases of WS were documented on Acroporids in 2017 while GAs were less host-specific found on nine genera across both years. The prevalence of compromised health was much higher than diseases with an average of 37.4 ± 3.9% across sites and years (Figure 5).

Prevalence of disease and compromised health categories varied significantly by year and site (repeated measures PERMANOVA, pseudo-F(3,47) = 3.7611; p = 0.0042) and site and depth interactions (pseudo-F(3,47) = 4.4228; p = 0.0094). Rural-N had the lowest prevalence of disease and compromised health compared to all sites in 2015 (22.43 ± 0.92%) and 2017 (33.84 ± 4.25%; p(perm) < 0.05). However, Rural-N was the only site where the prevalence of compromised health and disease increased between survey years. Despite this, Rural-N was characterized by the highest percentage of healthy corals (78.0 ± 0.9%), significantly higher than all other sites in 2015 but not significantly lower in 2017 (61.7 ± 4.7%; three-way ANOVA χ^2^(3,16)= 12.5135, p = 0.006; p < 0.05). This site also had the lowest prevalence of algal overgrowth on corals in 2015 (5.3 ± 1.2%) significantly lower than Urban-W in the same year (20.3 ± 1.8%; χ^2^(3,36) = 58.42713, p < 0.001) and the lowest amount of bleaching both years (6.0 ± 0.9% in 2015, 0.8 ± 0.2%; Table A4).

### 3.3 Water quality

#### 3.3.1 Nutrients and stable isotopes

Seawater nutrient levels and N stable isotopes of macroalgae were simultaneously assessed to get an indication of land-based pollution. Nutrients were not elevated at the urban sites where sewage pollution can result in > 10 μM DIN although there were significant site and depth interactions (two-way MANOVA F(3,63) = 3.208, Pillai = 0.398, p = 0.0012). Nitrate, nitrite, and phosphate were responsible for these interactions (two-way ANOVA 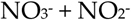: F(3,63) = 10.899, p < 0.001; 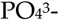: F(3,63) = 4.560, p = 0.006). Rural-N 10 m had significantly higher combined nitrate and nitrite (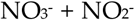: 1.05 ± 0.07 μM) and phosphate (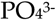0.15 ± 0.01 μM; Table A5), than all other sites at 10 m, but comparable levels of both at 5 m (Figure 8). DIN was marginally significant with a site and depth interaction (two-way ANOVA F(3,63) = 2.777, p = 0.0484), but pairwise test showed no significant comparisons (p < 0.05; Figure 6).

**Figure 6.**
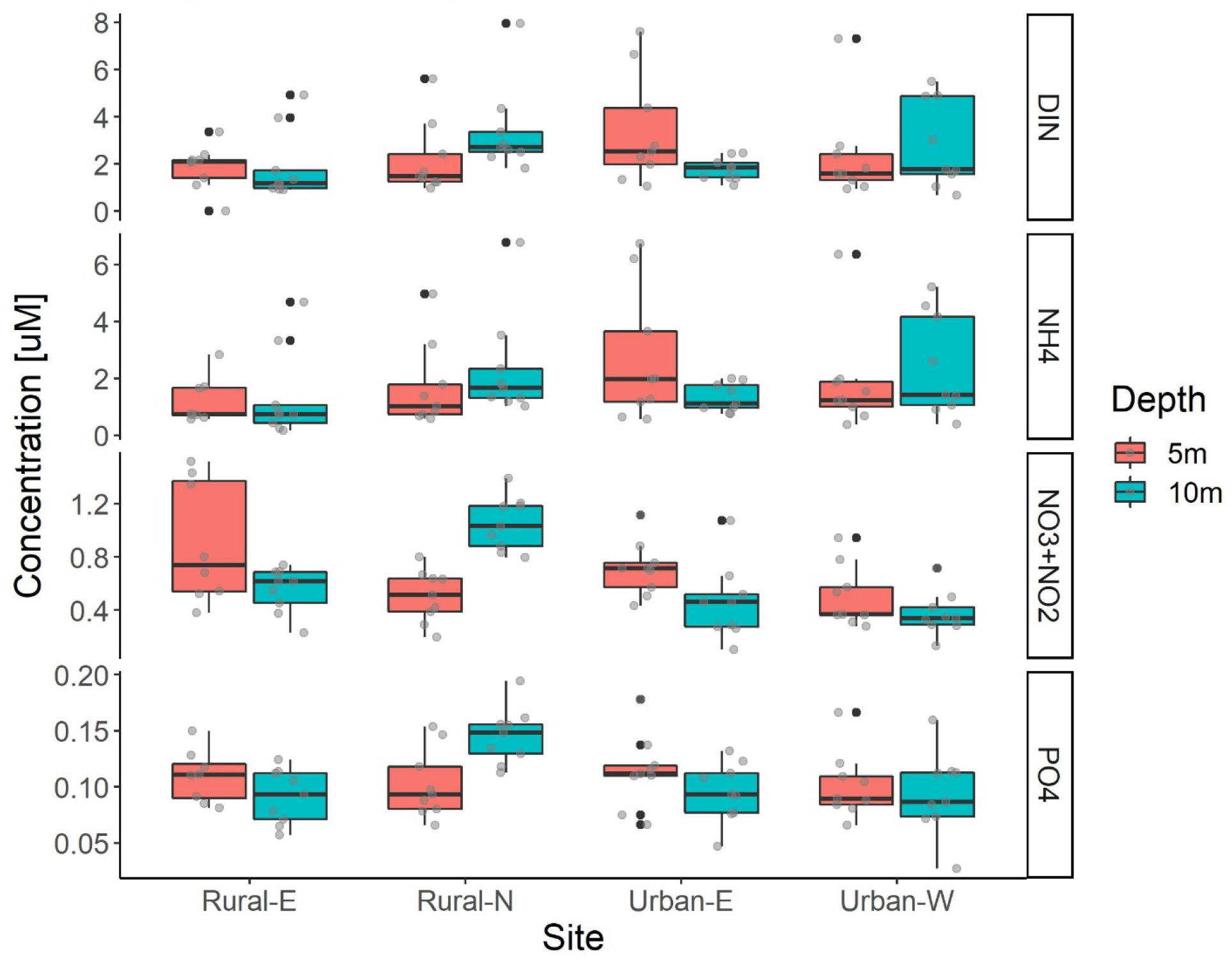
Seawater nutrient concentrations (top to bottom: DIN, 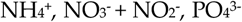) sampled in triplicate at four sites (Urban-W, Urban-E, Rural-N, Rural-E), two depths (5 m and 10 m), and three transects per depth in Timor-Leste in 2015. Bold line is the median, box ends are the first and third quartile, lines are 95% confidence interval of the median, and points are Tukey’s outliers.

Stable isotope values were remarkably consistent across sites, with no elevated levels at the urban sites compared to the rural sites. Delta ^15^N stable isotopes had a significant site difference for both algae species. Urban-E had significantly lower δ^15^N for both algae species (Table 2).

**Table 1.** Delta ^15^N stable isotope ANOVA results of two genera of algae sampled in replicates at the four sites (Urban-W, Urban-E, Rural-N, Rural-E), two depths (5 m and 10 m), and three transects per depth in Timor-Leste in 2015. Bolded values are significant results with mean, SE, and posthoc groupings presented per site. No *Chlorodesmis sp.* was sampled at Rural-N or Rural-E at 10 m and the three samples collected from a single transect at Rural-E 5 m were removed for the ANOVAs. N% and C:N ratio values and statistics are presented in Table A6.

| Algae | Effect | df | F-value | p-value | Rural-E | Rural-N | Urban-E | Urban-W |
| --- | --- | --- | --- | --- | --- | --- | --- | --- |
| Halimeda sp. | Site | 3 | 3.8199 | 0.0121* | 4.26‰ | 4.31‰ | 4.03‰ | 4.26‰ |
|  | Depth | 1 | 0.5442 | 0.4624 | ±0.01 | ±0.01 | ±0.01 | ±0.01 |
|  | Site x Depth | 3 | 1.3801 | 0.2531 | ab | b | a | b |
| Chlorodesmis sp. | Site | 1 | 10.0028 | 0.0064* | 4.57‰ | - | 4.11‰ | 4.47‰ |
|  | Depth | 1 | 0.1747 | 0.6819 | ±0.09 |  | ±0.02 | ±0.04 |
|  | Site x Depth | 1 | 2.4127 | 0.1412 |  |  | a | b |

Testing the drivers of prevalence of disease and compromised health with seawater nutrient concentrations in 2015, they differed significantly between site (p(perm) = 0.0001), depth (p(perm) = 0.0016), and 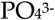(p(perm) = 0.0162; Table A7). The dispersion, or variability, grouped by site and depth was significantly different (one-way ANOVA F(7,16) = 5.931, p < 0.01) and dispersion was greatest at Rural-N and significantly larger than all other sites at 10 m and Urban-E and Rural-E at 5 m (p < 0.05). Phosphate was significantly higher at Rural-N 10 m compared to other sites at the same depth (Figure 6) and Rural-N had the highest prevalence of WS (Figure 5).

#### 3.4 Temperature and the prevalence of bleaching

The *in situ* logger temperature data was significantly different by a site and depth interaction (three-way ANOVA: F(2, 85) = 5.7503, p = 0.0045) and month nested in year (three-way ANOVA: F(15, 85) = 175.2521, p < 0.0001). Urban-E (29.37 ± 0.17°C) was not significantly different from the other two sites at 10 m and Rural-N (28.98 ± 0.17°C) and Urban-W (28.99 ± 0.17°C) were not significantly different at 5 m (p < 0.05). Additionally, mean monthly temperatures in 2016 were higher than the corresponding months in 2015 and 2017 in the six months where overlap occurred (Figure 7). Comparison of the monthly mean of all *in situ* loggers and CRWTL monthly temperature mean was significantly different by a season and site interaction (two-way ANOVA: F(3,53) = 3.92, p = 0.0100). The CRWTL 5 km satellite-derived SST was significantly higher than the *in situ* logger data during the austral summer (Jan–Mar, CRWTL = 30.67 ± 0.47°C, logger = 29.08 ± 0.50 °C).

**Figure 7.**
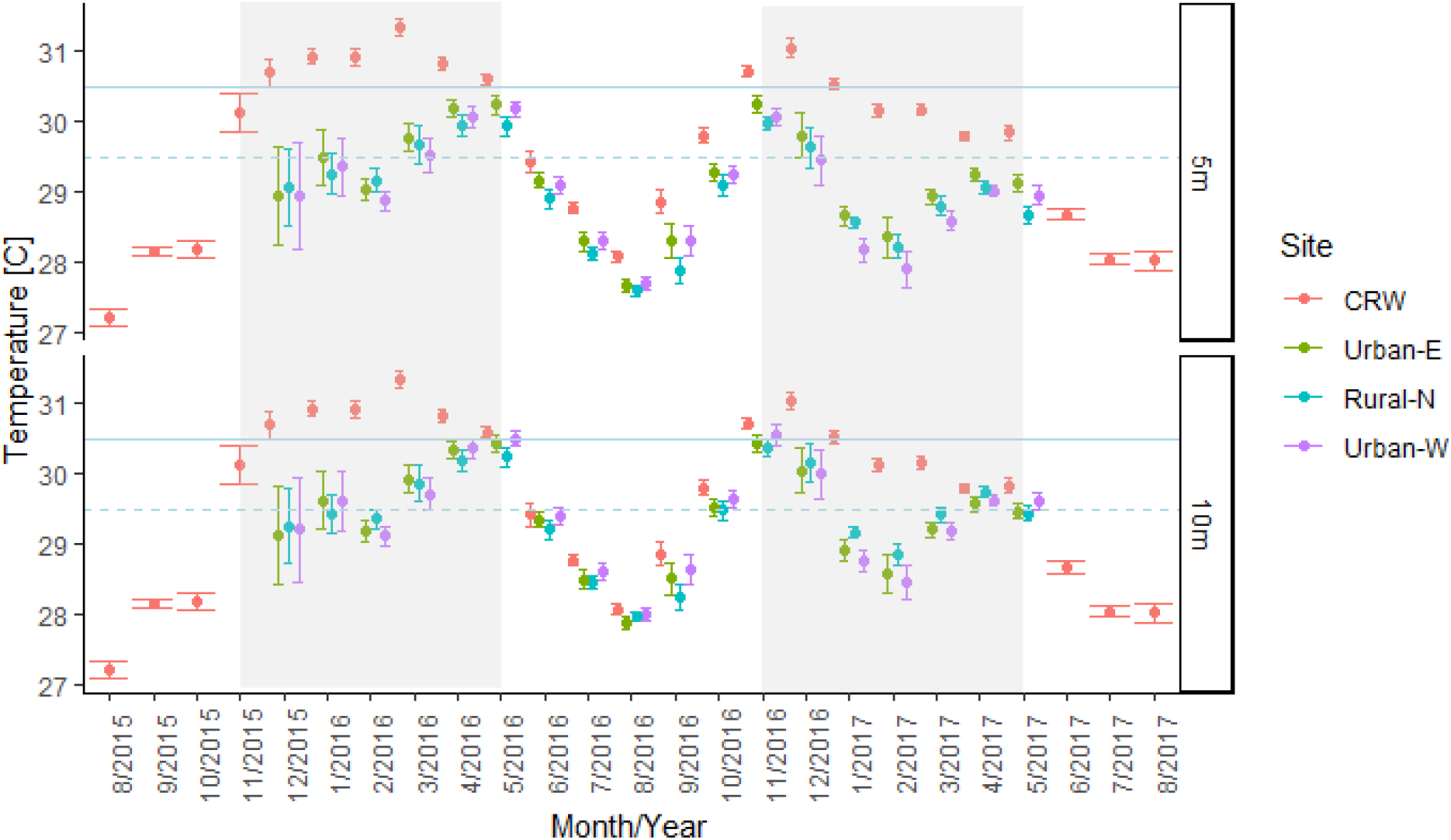
Mean temperature by month ± 2SE for remotely sensed sea surface temperature from Coral Reef Watch (CRW) and in situ temperature loggers (Rural-N, Urban-E, Urban-W at 5 and 10 m depth) between sampling periods in 2015 and 2017. The dashed blue line is the maximum monthly mean (MMM) from CRW data and the solid blue line is MMM + 1°C, the bleaching threshold.

A major heat stress event occurred between surveys in November 2015 and July 2017 (639 days). CRWTL indicated there was 190 days (30.2%) of bleaching warning (0 < DHW < 4) and 161 days (25.2%) at bleaching alert 1 (4 <= DHW < 8). The accumulation of DHWs was limited to November 29^th^, 2015–July 12^th^ of 2016 (224 days) and November 13^th^, 2016 through March 16^th^, 2017 (119 days) which corresponds to the months where the CRWTL monthly averaged temperatures were greater than the MMM (Figure 7). The accumulation of DHWs between 2015–2016 was almost 8 months, twice as long as the DHW accumulation from 2016–2017. The *in situ* temperature data, however, never reaches the MMM + 1°C threshold for bleaching, and based on these data there would be no accumulation of DHWs.

There was a three-way interaction between site, depth, and year on bleaching prevalence (repeated measures ANOVA, χ^2^ = 18.6709, p = 0.03; p < 0.05; Figure 5). All sites at the same depth showed a decrease in the prevalence of coral bleaching between surveys, which is expected as the second survey was conducted at the onset of austral winter. However, only Rural-E at 10 m (13.4 ± 0.7% and 2.8 ± .3% in 2015 and 2017 respectively) and Urban-W at 5 m (17.4 ± .6% and 1.8 ± 1.3%) had significant decreases. Rural-N in 2017 (1.1 ± .5%) had significantly less bleaching than all other sites other than Urban-E (3.3 ± 1.3%; p < 0.05). Paling also occurs with seasonal swings and photoacclimatory changes in symbiont density can be difficult to differentiate with bleaching in corals [89].

## 4. Discussion

In this study, we produced a baseline for the condition of four outer reef slope communities at rural and urban sites near the capital Dili, in Timor-Leste. A higher prevalence of coral disease and other signs of compromised health was expected at urban sites with elevated nutrients from sewage pollution. Our study reports major differences between sites in terms of community composition, disease prevalence, and potential exposure to local threats, but disease prevalence was low overall and nutrient levels were consistent across sites. Insights and answers were derived for two key questions posed at the outset of this study. Firstly, there is evidence of human subsistence activities influencing the health of a reef at one of the urban sites and second, Timorese reefs were subjected to bleaching during the 2016 global bleaching with the accumulation of >4 DHWs. This, however, was not supported by the *in situ* temperature logger data. Whilst recording similar temperatures in cooler months to that observed by satellite average for the region, the loggers recorded significantly lower temperatures over the summer months, never reaching the MMM + 1°C threshold associated with bleaching. Mortality associated with the event was low by comparison to regions that experienced > 8 DHW during the bleaching event and high coral mortality.

### 4.1 Coral community composition and human impacts

The underlying coral community composition was different across the four surveyed sites, which influenced the prevalence of coral disease. Rural-N had the highest coral cover and diversity dominated by Acroporids which is comparable to the biodiversity assessment of the same site in the 2012 Rapid Marine Assessment [90]. While damage to reefs may correlate with the local population density of humans [91–94], rural sites in the present study did not have a lower prevalence of coral disease. Although not definitive, there is some evidence suggesting the rural sites are in better shape in terms of coral cover than the urban sites. However, the marked presence of Acroporids at Rural-N seems to indicate that this reef is distinctive from the remaining sites versus clear rural and urban distinctions. Rural-N was the only barrier reef survey which are uncommon along the north coast and harder to access. It was anecdotally observed, that site-specific factors, such as ease of access to the reef, were associated with reduced reef health in terms of reduced coral cover, coral diversity, and other signs of compromised health.

Significant differences were identified between the state of coral reefs between Rural-N and the three mainland sites (Urban-W, Urban-E, and Rural-E). Localized impacts along the northern coast of Timor-Leste include watershed-based pollution and fishing and gleaning [15,20,95,96]. Geography, season, and factors such as land-use, accumulated wave exposure, and storm exposure are likely to be important but were not studied here. However, Rural-N may be subjected to less sedimentation than the other three sites, as Ataúro Island does not have any major rivers as on Timor island. Dili (encompassing Urban-E and Urban-W) and Rural-E in Manatuto are both near major rivers (the Comoro and Laclo rivers respectively). Large storms and waves are uncommon along the north coast. Temperature likely has a negligible influence on community composition as the temperature logger data was consistent between the three sites where loggers were successfully retrieved, Rural-N, Rural-E, and Urban-W (Figure 7). This leads us to localized human impacts as a key source of impact on coral reefs.

Fishing is playing an increasingly significant role in Timor-Leste. Observations of extensive rubble slopes at Urban-W may be due to blast fishing although the damage does not appear to be recent [90]. Gleaning is largely overlooked when assessing fisheries in Southeast Asia although most (> 80%) of households in coastal communities participate in gleaning activities in Timor-Leste [97,98]. Women and children glean while men fish, however, gleaners have nearly a 100% success rate highlighting its importance in maintaining food security especially during low crop periods [97,99]. Increased gleaning could also be a sign of diminishing fishing returns [100] or economic crises [101]. Gleaning may have played an even greater role in food security during recent times of violence and instability resulting in degraded coral reef flats, particularly in densely populated areas. Human gleaning and the associated trampling of intertidal reefs have been demonstrated to have deleterious effects on coral cover on reef flats although depths greater than 5 m are generally out of reach [15,102,103].

Urban-W site had more fishing activity than the other sites from observations while conducting fieldwork and also had the greatest signs of blast fishing. The subdistrict of Dom Alexio encompassing this site had the highest population density out of the four sites with 5,017.9 people/km^2^ compared to 779.5 people/km^2^ at Urban-E and 79.3 people/km^2^ nationally. The low coral cover at 5 m and the minimal diversity at 10 m could be attributed to the high subsistence and recreational usage at this site. Women were gleaning for invertebrates on the low tide, small children were playing in the surf and on the reef flat, and men were net-fishing from small boats (Figure A3). While distance to river is a sensible explanation between community-level differences between Rural-N and Rural-E, relative ease of access in a densely populated area differentiated Urban-W. Urban-W is in walking distance to a densely populated area with fishing boats lining the beach while Urban-E is tucked in at the edge of Dili Bay surrounded by steep hills and more affluent neighborhoods.

### 4.2 The health of coral reefs along the north coast of Timor-Leste

Prevalence of disease and compromised coral health was expected to be greatest at urban sites with larger nutrient input and greater δ^15^N values at the shallow 5 m surveys. Contrary to expectations, disease was highest at Rural-N at 5 m with levels of WS at ~1% in both survey years. Combined nitrate and nitrite, and phosphate levels were also highest at Rural-N, but at 10 m. The low levels of disease detected in the current study agree with previous surveys [90,104], although no previous studies were specifically quantifying disease and compromised health.

WS was the main pathology consistently observed during surveys. The WS documented at Rural-N was likely an infectious disease [25] and in the Indo-Pacific, WS is known to target Acroporids [7,25,105]. Direct transmission of WS spreading between Acroporid corals was observed in the field, in addition to a positive association between host abundance and disease prevalence. This follows the classic density dependent host pathogen relationship [90–92], a phenomenon which has been demonstrated in coral disease ecology. In this study, all but one case of WS was on Acroporids. In 2015, 13 of the 17 recorded cases were at Rural-N, while in 2017 all 10 cases were documented at Rural-N which had the highest density of Acroporids [7,23,106,107]. The few cases at other sites could have been from other causes such as unidentified predation; however, there was sufficient evidence to suggest that WS at Rural-N was caused by an infectious pathogen.

There was likely coral mortality caused by the WS, inferred from the proportion of dead coral on some colonies (Figure 2a) as is typical with WS progression on Acroporids [108,109]. While a low prevalence of WS was at Rural-N 5m it is likely not responsible for the decrease in coral cover. Prevalence of WS did not differ between depths (Figure 5), and there was an increase in coral cover at Rural-N 10 m. WS recorded here is likely typical background levels of disease comparable to other CT locations versus an outbreak although the prevalence of WS should continue to be monitored [107,110–113]. The low prevalence of coral disease in the CT supports the disease-diversity hypothesis which predicts that higher host species diversity should result in decreased severity of a specialist pathogen through increasing interspecific competition [114–116]. The majority (> 50%) of cases were on tabulate Acroporids and the most susceptible as documented on other Pacific reefs to WS out of four morphologies (likely different species) identified during surveys [114].

WS is a dynamic disease and can occur in outbreaks devastating Acroporid populations [25,117] and thus alter overall coral community structure [117]. The pathogen causing WS at Rural-N is unknown but was likely a *Vibrio spp.* bacteria that have been associated with diseases of multiple marine organisms including corals and humans [118–121]. These bacteria have previously been implicated as a causative agent of WS [122–124]. The causes of WS outbreaks have been linked to sediment plumes from dredging and terrestrial runoff and elevated ocean temperature [7,8,125,126]. This is especially relevant given the recent global bleaching event and expected increase in the prevalence and severity of marine diseases given continued ocean warming [127]. A significant relationship between WS and coral bleaching co-infection was found on the GBR during the 2016– 2017 global bleaching event. Acropora colonies that exhibited both WS and bleaching had seven times more tissue loss than solely bleached colonies [75]. Cooler temperatures could be a protective factor against outbreaks of WS in Timor-Leste given the cooler subsurface temperatures on reefs compared to SST during the monsoon season which coincides with the yearly ocean temperature maximum. Increased sedimentation from catchments, however, is a continued threat as watershed health in Timor-Leste is poor and there is a need for future work assessing impacts of sedimentation on reefs and coral health.

The prevalence of indicators of compromised health was much greater than the prevalence of disease at surveyed sites. Rural-N at 5 m had the highest prevalence of non-coral invertebrate overgrowth (Figure 5), which could be explained by greater coral cover eliciting more coral-invertebrate interactions as the cover of invertebrates is comparable between all sites. The infestation of flatworms was found at all sites except Urban-W 10 m with similar prevalence as surveys in Indonesia [107] with some severe cases (Figure 2c). Flatworms consume coral mucus, reduce heterotrophic feeding, and decrease photosynthesis in high densities [128–130] although their role in coral reef environments is not well understood. There was also a notable absence of turf overgrowth at Rural-N while the remaining sites have high levels which could be indicative of depauperate herbivore communities or elevated nutrients at these locations [131,132]. Different competitive interactions such as burrowing barnacles, CCA overgrowth, and turf overgrowth were also more commonly found on genera with specific morphologies significantly associated with *Platygyra, Montastrea*, and massive *Porites*, all massive species. The highest cyanobacteria overgrowth was found at Rural-W at 10 m which was most closely associated with branching Montiporids and Poritids.

### 4.3 Water quality andsources of nutrients in Timor-Leste

The nutrient concentrations plus stable isotope ratios were more indicative of oceanic processes (i.e., upwelling, internal waves, etc.) versus terrestrially based nutrient pollution. Phosphate was a significant driver of the prevalence of disease and compromised health (Table A7) and the highest values of each parameter were both at Rural-N which suggests that increased phosphate is associated with a higher prevalence of disease (Figure 6; Figure 5). High levels of inorganic nutrients are a major driver of reef degradation [28] although there has been debate on whether nutrients or overfishing are more important [6,133]. The levels of inorganic nutrients found in this study were comparable to the low values of nutrients measured in the Laclo river, near Rural-E, in 2006 (See Appendix) [20] and not indicative of nutrient pollution.

Contrary to expected, combined nitrate and nitrite and phosphate averages were highest at Rural-N at 10 m which could be indicative of upwelling [64,134,135]. Slightly elevated nutrients off Ataúro Island in the channel at 10 m may suggest upwelling of deeper, nutrient-rich water [90,136]. Other sources of nutrients at depth to consider is submarine groundwater discharge [137]. Cyanobacteria overgrowth was only found at high levels at Urban-W 10 m in 2015 (6.1% prevalence) which can be a sign of elevated nutrients or other disturbances (e.g., ship strikes, etc.) [138–140]; however, seawater nutrients and stable isotope values were not significantly higher at this site during our sampling. In 2017, the prevalence of cyanobacteria decreased to 0.0% indicating cyanobacteria in 2015 was an ephemeral bloom. Although, 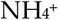 was not significantly different between surveyed sites except Rural-E 5 m, the range of 1.32 ± 0.17 μM to 2.69 ± 0.78 μM was more than the previously recorded values between 0.3 μM and 2.2 μM on reefs [49,141,142].

The stable isotope data were consistent across sites and depths sampled (range 2.5–5.5‰ excluding outliers) and fall within pristine oceanic values of 2–3‰ [44,143] and upwelling values of 5–6‰ [46,59,61–63]. Sewage-affected waters have generally higher δ^15^N values from 8–22‰ [45,48,57,144,145] and our data was not indicative of high δ^15^N enrichment. Although the mean was significantly higher for the *Chlorodesmis sp.* at Urban-W versus Urban-E, the sampling of that alga was sparse compared to the *Halimeda sp.* Additionally, there was no *Chlorodesmis sp.* found at Rural-N. Calcareous algae are good integrators of nitrogen over weeks to months versus days with fleshy macroalgae [56]. Similar values were recorded for both algae collected across sites and depths which indicates that the influx of nitrogen has been stable across several months. This is likely due to sampling at the end of the dry season (Mar to Nov) with little terrestrial runoff. There were a few outliers of much higher (12.17‰, 15.12‰) and lower (−6.79‰) δ^15^N values within *Halimeda sp.* which could be indicative of localized inputs on a scale of tens of meters of nutrients such as fish waste or groundwater discharge. Previous studies reveal that macroalgal δ^15^N signatures decrease with depth on range from 5 to 35 m because of land-based pollution [46,48,53,146]. The influence of upwelling is less clear as both δ^15^N depletion and enrichment have been reported with upwelling [46,61,64,147].

In summary, assigning direct links between the condition of coral reefs and the source of nutrients is difficult. In the present study, the mean δ^15^N values of 4.3‰ and 4.2‰ for *Chlorodesmis sp.* and *Halimeda sp.* respectively are higher than those reported from the open ocean. Given that the algae were collected at the end of the six-month dry season, it is unlikely our sampling captured the effects of terrestrial and river run-off and potential sewage pollution. Significant seasonal differences in sampling of macroalgae for stable isotope analysis have been demonstrated [47] and further seasonal investigations could elucidate the source of nutrients in nearshore waters. The seawater nutrient and stable isotope data are likely indicative of oceanic influence with potential upwelling in the absence of aquaculture industries and heavy use of inorganic fertilizers and pesticides in-country coupled with sampling conducted after months of no rain. The higher seawater ammonium, nitrate, nitrite, and phosphate nutrients at Rural-N 10 m depth could indicate short upwelling events that are too ephemeral to be assimilated by calcareous macroalgae over weeks to months. Rural-N’s location in the Timor Strait and ITF with the large volumes of water movement through the channel [148] could be more conducive to localized upwelling than the mainland sites.

### 4.4 Elevated temperature and the prevalence of bleaching from thermal stress

The surveys in the present study were conducted right before the austral summer during the 2015 ENSO event which triggered mass bleaching worldwide [21]. The CRWTL virtual monitoring station indicated that the temperature began rising above the monthly maximum mean in November 2015; however, care must be taken given that the satellite data only measures the temperature of the first 10–20 μm of the ocean surface [149]. Satellite temperature products can therefore be ineffective in coastal waters due to pixels mostly encompassing open ocean versus coastal waters. Additionally, Timorese reefs are very steep and close to the coast [136] where satellite data are unreliable due to the potential interference of land temperatures.

*In situ* temperature varied between the three sites and between months. Most interestingly, there was a divergence between the *in situ* and CRWTL temperature data during the austral summer months. The surveys in 2015 were conducted in November at the onset of austral summer and the yearly ocean temperature maximum. In 2017, the surveys were conducted in July approaching the yearly ocean temperature minimum. Timor-Leste appeared to have experienced lower levels of bleaching compared to other regions of the world such as the Northern Great Barrier Reef (NGBR), one of the most severely affected by bleaching. The CRWTL accumulated DHWs for 55% of the days between survey periods compared to 49% of days over the same time in the NGBR. The magnitude of DHWs in the NGBR reached 13.59°C-weeks, more than double the 5.79°C-weeks maximum in Timor-Leste. Comparison of *in situ* bleaching surveys and DHWs on the GBR indicate that 2-3°C-weeks are associated with low levels of bleaching, > 4 °C-weeks with 30-40% corals bleached, and > 8 °C-weeks with an average of 70-90% of corals bleached [67,68]. The bleaching severity of the NGBR was greater than 60% bleached for all surveyed reefs in 2016 and, although there is no data on the extent or severity of bleaching on reefs in Timor-Leste, DHW data would project mass coral bleaching in Timor-Leste of around 30–40%.

Local dive operators reported mass coral bleaching at Jaco Island at the end of March. By the end of May, 90% of *Goniopora sp.* on Ataúro Island were bleaching (Figure A4a), massive *Porites sp.* from 5–18 m at Jaco Island (Figure A4b), and staghorn Acroporids in the shallows of the same area. Bleaching reportedly began in the shallows and progressively affected corals at deeper sites. The timing of the bleaching also matches the *in situ* temperature logger data where the mean monthly temperatures exceeded the MMM in March 2016 versus November 2015 for the CRWTL SST data (Figure 7). The *in situ* temperatures never exceeded the MMM + 1°C threshold for mass bleaching. The range of the temperature loggers during December 2015 was from 27°C to almost 31°C so reefs did experience elevated temperatures, but not for prolonged periods. The *in situ* mean temperature began to creep over the MMM and close the gap with the CRWTL data in March and April of 2016. The loggers approached MMM + 1°C in May of 2016 five months after the CRWTL temperatures had been above the bleaching threshold (Figure 7). The *in situ* data is limited to the Dili, Ataúro Island, and Manatuto areas which may not be representative of temperature regimes in the Jaco Island region. Even so, anecdotal evidence of the most bleaching in May 2016 in both Ataúro and Jaco Islands follows the temperature timeline of the logger data. CRWTL predicting bleaching too early in Timor-Leste.

Based on the comparison of *in situ* temperature logger data in Timor-Leste and the satellite-derived SST, CRWTL overestimates the bleaching stress in-country. During the austral summer CRWTL SST was more than 1.5°C warmer than the *in situ* data at 10 and even 5 m depth in both years (Figure 7). This could be due to seasonal changes in the oceanography that increased water movement, or increased upwelling along the north coast coinciding with the northwest monsoon during the austral summer. The significant divergence between the *in situ* and CRWTL temperatures, during this time of year, point to seasonal oceanography (i.e., upwelling or internal waves) influencing temperature on shallow reefs without reaching the surface to affect remotely sensed SST. The northwest monsoon season from December to March is associated with a weak reversal in the flow of the ITF [150]. Whether this is associated with coastal upwelling remains unclear; however, there is clear seasonal variability of the ITF. Additionally, the temperature range in December 2016 is 27–31°C, nearly 4°C, in all six loggers compared to 29–31°C in December 2017. ENSO could be strengthening upwelling along the north coast of Timor-Leste bringing up cooler water to shallow reefs while the SSTs are elevated above the bleaching threshold.

The effect of upwelling could be a protective factor for Timorese reefs against climate change; however, the divergence in temperatures between CRWTL and logger temperature data appears to be seasonal as the temperatures converge around April/May of 2016 when seasonal upwelling may subside (Figure 7). Coral bleaching did occur in Timor-Leste only not to the same extent as other reef regions. Although this mitigation of elevated temperature is positive, other potential impacts must also be considered. For example, the exposure to cooler upwelled waters regularly acclimatizes the corals to these conditions and could potentially make them more sensitive to elevated temperatures once upwelling stops. Additionally, Timor-Leste was identified as having the lowest calcification rates out of all the NOAA Indo-Pacific monitoring sites [79] which could be associated with hypercapnic (CO_2_-rich) upwelled waters [151]. This lower calcification could affect Timorese reef’s ability to cope with sea-level rise and recover from disturbances such as physical damage and bleaching. There is, however, evidence that calcifying organisms are able to withstand seasonal increases in acidity potentially relying on increased heterotrophic feeding [152,153]. Clearly, further research on the oceanography of the region and interactions between environmental parameters (light, temperature, CO_2_, salinity, etc.,) are critical to understanding and effectively managing the country’s marine resources.

## 5. Conclusion

The present study set out to understand the nature of both local and global threats to the relatively understudied coral reefs of Timor-Leste. Baseline information on these systems is limited despite the current and future importance of these marine resources to Timor-Leste. There were two major objectives of the present project. The first was to explore the state of coral reefs, particularly concerning the presence of coral disease or other evidence of compromised health and benthic composition. The second was to assess the impact of humans on issues such as water quality and climate change. Coral reefs of the north coast of Timor-Leste are characterized by high amounts of coral cover as much as 58.2 ± 6.4%. The concern, however, is that sites close to the urban areas of the capital city, Dili, are showing signs of significant degradation with < 5% hard coral cover at 5 m at one of the two urban sites. Also, there is the global problem of heat stress from climate change driving additional pressure on coral reef systems. Like coral reefs everywhere, a failure to act to drastically and reduce emissions of greenhouse gases from burning fossil fuels and land-use change will see further and more rapid degradation of the coral reef resources of Timor-Leste [154].

The human population density of sites did not directly correlate with the condition of coral reefs. The only consistent presence of infectious coral disease was found only at a rural site with low human population density. Other signs of compromised coral health such as algal overgrowth, burrowing barnacles, etc. were much more widespread across all sites. The underlying differences in coral community structure were key to the prevalence of WS on tabulate Acroporids which may have been shaped by human impacts such as subsistence livelihoods and degradation of watersheds. There were two lines of evidence at local scales, temperature, and nutrients (both seawater concentrations and algal δ^15^N signatures), that indicated upwelling is a significant feature influencing the shallow reefs of the north coast of Timor-Leste. This upwelling appears to lessen the impact and length of bleaching events, although more research across larger spatial and temporal scales is necessary. While it is fortunate the mass bleaching event did not cause large scale coral mortality in Timor-Leste, the sublethal effects pose another threat to already highly impacted reefs.

Over 300 million people rely on coastal resources for economic livelihoods and cultural practices in the CT [155]. Sustainable management of these resources ties into larger socio-economic issues such as food security in Timor-Leste. Fishing is an important means of protein and nutrients and vital for food security nationally and dominated by artisanal, low-efficiency methods [17,155,156]. The fishing industry in-country is largely contained to subsistence practices because of a lack of reliable road, electric, and refrigeration infrastructure. As these systems are developed and the capabilities and supply chains to transport fish increases and these developments must be monitored. Additionally, *meti*, or gleaning, is an important component of subsistence fishing. In urban areas, the differences between ease of access between Urban-W and Urban-E seemed to play an important role in the types of human impacts present (i.e., recreational versus extractive). Localized impacts such as fishing (including blast fishing pre-independence) and gleaning, which over the last several decades, were observed and likely affected coral cover and diversity in both urban and rural areas.

To address these large and small-scale impacts, coral reef management needs targeted local and national actions. Ultimately, the health of coral reefs is tied to the unique social and economic development needs of the country as a whole. At a local level, the resurgence of t*ara bandu* with increased interest in community marine protected areas (MPAs) is promising for the future of Timorese reefs. *Tara bandu* is customary law that administers prohibition designations in communities banning practices such as tree-cutting [157–159]. These designations are well-adhered to within communities, but appear to be more effective in rural areas [158]. For MPAs, certain reefs are established as no-take, ecotourism zones where visitors pay a small fee for snorkeling and diving. The reef housing the Rural-N site in this study was designated as an MPA six weeks before the resurvey in July 2017 and a small fee was paid. Fishermen were witnessed bypassing the reef by canoe to fish on the adjacent reef to the north. Currently, the formation of MPAs is centered on Ataúro Island, but the success and income generated for the community are seeing this practice expand to other communities. The level of community engagement in designating marine reserves provides a positive outlook for coral reef management although the long-term impacts of COVID-19 remain to be seen.

Assuming the international community takes strong action under the Paris Climate Agreement, it will be very important for countries such as Timor-Leste to establish effective management of its coral reef resources. This lies at the heart of international strategies such as the 50 Reefs project and the evidence here corroborates Timor-Leste’s inclusion as a climate robust reef region [160,161]. At a country-scale, developing a national network of MPAs, rebuilding healthy watersheds, and understanding vulnerability to climate change is key to the sustainable management of coral reefs. Ideally, a systematic approach would be taken to establish no-take and mixed-used zones across all Timorese waters incorporating MPAs and the national marine park. Establishing a national plan before the development of tourism and industrial industries would be beneficial in ensuring a certain level of sustainable management of coastal resources. Lastly, monitoring the heat stress and bleaching of Timorese reefs must continue and further research of the oceanography along the coasts (especially possible benefits from upwelling) will advance the understanding of the country’s vulnerability to climate change as a whole.

## Acknowledgments

We gratefully acknowledge the following sources of funding that supported this research: the Global Change Institute at the University of Queensland, the Society of Conservation Biology, and the Winifred Violett Scott Trust. Many thanks to the Ministry of Agriculture and Fisheries in Timor-Leste, Trudiann Dale and the marine team at Conservation International Timor-Leste, volunteers Dominic Bryant, Craig Heatherington, Peran Bray, and Fiona Ryan for fieldwork support, Sarah Naylor, Rachel Hazel, Abbie Taylor, Shari Stepanoff, and Susie Green for logistical support. Thank you to the Compass Boating & Diving and Aquatica crews. Samples were exported from Timor-Leste under export permit No. 455 and imported to Australia under AQIS import permit IP15000663.

## Author Contributions

Conceptualization, C.K. and S.D.; methodology, C.K. and S.D.; formal analysis, C.K. and S.D.; investigation, C.K..; resources, C.K.; data curation, C.K.; writing—original draft preparation, C.K.; writing—review and editing, C.K., S.D., O.H-G.; visualization, C.K.; supervision, S.D., C.R., and O.H-G.; project administration, C.K.; funding acquisition, C.K. and O.H-G. All authors have read and agreed to the published version of the manuscript.

