## Supplemental Materials for "The condition of coral reefs in Timor-Leste before and after the 2016–2017 marine heatwave"

**Supplementary Materials:**

Table A1 An overview of coral disease prevalence within the Coral Triangle. Disease abbreviations are as follows: AtrN–Atramentous necrosis, BrB–Brown band disease, BBD–Black band disease, GAs–Growth anomalies, N–Necrosis, SATL–Subacute tissue loss, SEB–Skeletal eroding band, PR–Pigmentation Response, PUWS–*Porites* Ulcerative White Spot, UWS–Ulcerative white spot, WP–White Plague, WS–White syndrome, YBD–Yellow band disease

| Location | Diseases | Prevalence | Reference |
| --- | --- | --- | --- |
| Included: Bali, Sulawesi, West Papua, Indonesia | BBD, BrB, GAs, SEB, WS | Range: 0.1 ±0.09% – 43.9 ±5.1% | [162] |
| Spermonde Archipelago, Indonesia | BBD, BrB GAs, SEB, WS | Total disease: 0.2 ±0.1%, 5.9 ±1.1% | [163] |
| Kepulauan Seribu Marine National Park, Indonesia | BBD, WS | <1% of each | [111] |
| Panjang Island, Indonesia | UWS, WP, YBD | WP: 17.76 ±8.6%; UWS: 6.59 ±0.08%; YBD: 2.88 ±0.05% | [164] |
| Spermonde Archipelago, Indonesia | AtrN, BrB, GAs SEB, WS | Shallow: 8.2 ±9.9% Deep: 9.6 ±2.7% | [112] |
| Wakatobi National Park, Indonesia | GAs, WS | 2007: 0.6% 2005: 0.3% | [106,165] |
| Tubbataha Natural Marine Park, Philippines | BBD, BrB, GAs, SEB, UWS, WS | Total disease: 2.4 ±0.1% – 13.5 ±5.4% | [166] |
| Cebu, Philippines | GA, PR, SATL, WS | Fished: 2.26 ±2.27% MPA: 7.13 ±3.41% | [167] |
| Central Philippines | BBD, BrB, GAs, SEB, UWS, WS | Range: 0.25 – 7.9% | [168] |
| Philippines | *Porites* GAs, PUWS | PUWS: 0 – 53.7% GAs: 0 – 39.1% | [169] |
| Central Visayas, Lingayen Gulf, Philippines | GA, N, PR, PUWS, WS | Total disease mean: 8.3 ± 1.2% | [170] |
| Sumilon Island, Philippines | PUWS | Range: 0 – 72.6 ±16.5% | [171] |
| Sabah, Borneo, Malaysia | AtrN, BBD, BrB, PR, WS | 0 – 0.44 colonies per $m^2$ | [172] |

Table A2 Summary of Indo-Pacific studies assessing nutrients on reefs. DIN–dissolved inorganic nitrogen, TN–total nitrogen, TP–total phosphorus

| Location | | Nutrient | Concentration [μM] | Reference |
|---|---|---|---|---|
| west Hawai'i Island | reef slope 12 m | $NH_4^+$ | 0.716 ± 0.22 | [39] |
| | | $NO_3^-$ | 0.38 ± 0.11 | |
| | | $PO_4^{3-}$ | 0.24 ± 0.07 | |
| | enriched | $NH_4^+$ | 148.80 ± 7.51 | |
| | | $NO_3^-$ | 94.35 ± 19.68 | |
| | | $PO_4^{3-}$ | 22.38 ± 2.06 | |
| Florida Keys | 5-6 m | DIN | 1.15 ± 0.05 | [40] |
| | enriched | DIN | 3.91 ± 1.34 | |
| Kingman Atoll | 10-12 m depth | DIN | 1.3 ± 0.08 | [37] |
| | | $PO_4^{3-}$ | 0.1 ± 0.003 | |
| Kiribati Atoll | | DIN | 3.6 ± 0.1 | |
| | | $PO_4^{3-}$ | 0.3 ± 0.024 | |
| Maui | surface 0.25 m | DIN | 0.1 ± 0.1-25.6 ± 17.8 | [36] |
| | | $PO_4^{3-}$ | 0.06 ± 0.06-0.43 ± 0.86 | |
| | coastal groundwater | DIN | 1.6 ± 1.8-414.9 ± 37.8 | |
| | | $PO_4^{3-}$ | 1.01 ± 1.03-4.90 ± 1.06 | |
| Philippines | 1-3 m | TN | 5.9-15 | [169] |
| | | TP | 0.13-0.98 | |
| Majuro Atoll | reef flat | $NH_4^+$ | 0.34 ± 0.15 | [38] |
| | | $NO_3^-$ | 0.45 ± 0.45 | |
| | | $PO_4^{3-}$ | 0.34 ± 0.15 | |
| | ground water wells | $NH_4^+$ | 2.4-16.9 | |
| | | $NO_3^-$ | 49-1,060 | |
| | | $PO_4^{3-}$ | 0.2-22.1 | |

Table A3 Description of coral diseases and compromised health found during surveys in Timor-Leste and references citing negative impacts to corals. Corresponds to figures 2 and A1.

| Disease/Condition | Description |
|---|---|
| **White Syndrome** | A class of tissue loss disease producing white symptoms in the Caribbean and the Indo-Pacific have led to the general description White Syndrome (WS) describing corals with clearly defined lesion of exposed coral skeleton from the rapid sloughing of coral tissue ranging from 1.0 to 124.6 cm of tissue loss per day not associated with predation (i.e., *Drupella* sp. snails or crown-of-thorns seastars-COTS). In the Indo-Pacific, WS predominantly affects tabulate Acroporids [7,25,173,174]. |
| **Growth Anomalies** | Gross lesions of raised tissue with lighter pigmentation and enlarged variable polyp found on several coral genera [175]. Energetically relies on resources from surrounding healthy tissue [176]. |
| **Trematodiasis** | Pink, swollen nodules caused by infection of a larval trematode in Porites corals which can reduce colony growth [177,178]. |
| **Burrowing Invertebrates** | Worms, barnacles that burrow in or on the surface of corals. Vermetids worms have been shown to change morphology and reduce growth [179] and burrowers have been associated with reduced skeletal strength [180]. |
| **Flatworm Infestation** | An infestation of flatworms, typically *Waminoa sp.*, on the surface of corals. Have been associated with tissue loss [181], reduced heterotrophic feeding [130], light-shading [128,182], and experimentally shown to feed on coral mucus [129]. |
| **Predation** | Scars or tissue loss from predation by *Drupella sp.* snails, COTS, and reef fishes [183]. Typically, invertebrate predators (Drupella, COTS) will be visible near the predation scar and reef fishes make distinctive lesions. |
| **Tissue Loss** | Unexplained tissue loss (absence of predator, shape of lesion, etc.) and possibly cause by infectious agents, physiologic disorders, or toxins [183,184]. |
| **CCA overgrowth** | CCA in competition with living coral causing a reaction or overgrowth. Two species of CCA have been identified to overgrow live coral [185,186] in the Indo-Pacific and one species in Yemen [187]. |
| **Cyanobacteria and turf overgrowth** | Cyanobacterial filaments or filamentous turf algae competing with or growing on living coral tissue. Cyanobacteria are often associated with eutrophication and can smother corals [188] and reduce recruitment [189]. Decreased coral function such as decreased zooxanthellae density and tissue thickness have been associated with turf-coral interactions [190]. |
| **Sponge or tunicate Overgrowth** | Sponge overgrowth on live coral tissue can negatively affect corals [191] with some aggressive species such as *Terpios sp.* and *Cliona sp.* killing corals [192]. Colonial tunicates smothering live coral [193–195]. |
| **Pigmentation response** | Pigmented edge of a coral lesion, may be swollen, form bumps, or irregular shapes. May be caused by borers, competitors, breakage, cyanobacteria [196], polychaetes, molluscs [25]. |

Table A4 Average (± SE) percent coral cover, diseased corals, and corals exhibiting other signs of compromised health, average number of genera, density of hard corals (colonies/$m^2$), total number of colonies surveyed per site and depth. % D–percent disease, % Comp–percent compromised health, # Gen–number of genera, Total # Col–total number of colonies

| Site | % Coral | % Disease | % Comp | # Gen | Density col/m2 | Total # Col |
|---|---|---|---|---|---|---|
| Rural-N 5m | 58.18 ± 1 .74 | 1.88 ± 0.18 | 13.56 ± 1.00 | 29.6 ± 4.0 | 8.4 | 672 |
| 10m | 42.54 ± 5.82 | 1.52 ± 0.77 | 16.02 ± 2.44 | 31.6 ± 1.5 | 7.4 | 759 |
| Rural-E 5m | 20.17 ± 3.49 | 0.21 ± 0.21 | 42.58 ± 3.64 | 28.6 ± 2.1 | 6.0 | 547 |
| 10m | 22.10 ± 2.59 | 0.47 ± 0.24 | 25.95 ± 5.78 | 26.3 ± 3.5 | 5.5 | 502 |
| Urban-W 5m | 4.80 ± 1.78 | 0.71 ± 0.71 | 33.48 ± 4.18 | 24.6 ± 4.2 | 3.8 | 350 |
| 10m | 45.23 ± 8.49 | 0.38 ± 0.38 | 37.44 ± 4.87 | 17.6 ± 4.0 | 6.4 | 581 |
| Urban-E 5m | 24.43 ± 6.35 | 0.42 ± 0.42 | 27.71 ± 5.98 | 27.3 ± 2.5 | 7.1 | 647 |
| 10 m | 12.29 ± 3.24 | 0.23 ± 0.23 | 28.77 ± 1.35 | 25.3 ± 1.5 | 5.4 | 492 |

Table A5 Average seawater nutrient values from samples collected in triplicate from transects at four sites in Timor-Leste in 2015. DIN is the sum of $NH_4^+$, $NO_2^-$, and $NO_3^-$. All units are in µM.

| Site | Depth | DIN | NH4+ | NO2-, and NO3- | PO4+ |
|---|---|---|---|---|---|
| Rural-E | 5m | 1.87 ± 0.31 | 0.00 ± 0.00 | 0.80 ± 0.17 | 0.00 ± 0.00 |
| Rural-E | 10m | 1.9 ± 0.49 | 1.35 ± 0.52 | 0.55 ± 0.06 | 0.09 ± 0.00 |
| Rural-N | 5m | 2.19 ± 0.51 | 1.69 ± 0.49 | 0.51 ± 0.07 | 0.10 ± 0.01 |
| Rural-N | 10m | 3.39 ± 0.62 | 2.34 ± 0.61 | 1.05 ± 0.07 | 0.15 ± 0.01 |
| Urban-E | 5m | 3.4 ± 0.78 | 2.69 ± 0.78 | 0.71 ± 0.07 | 0.11 ± 0.00 |
| Urban-E | 10m | 1.78 ± 0.16 | 1.32 ± 0.17 | 0.46 ± 0.09 | 0.10 ± 0.00 |
| Urban-W | 5m | 2.31 ± 0.66 | 1.81 ± 0.60 | 0.50 ± 0.08 | 0.10 ± 0.01 |
| Urban-W | 10m | 2.79 ± 0.62 | 2.41 ± 0.60 | 0.37 ± 0.05 | 0.09 ± 0.00 |

Table A6 N% and C:N ratio ANOVA results of two genera of algae sampled in replicates at the four sites (Urban-W, Urban-E, Rural-N, Rural-E), two depths (5 m and 10 m), and three transects per depth in Timor-Leste in 2015. Bolded values are significant results with mean, SE, and posthoc groupings presented per site. No *Chlorodesmis sp.* was sampled at Rural-N or Rural-E at 10 m and the three samples collected from a single transect at Rural-E 5 m were removed for the ANOVAs.

| Halimeda sp. | | df | F-value | p-value | | Rural-E | Rural-N | Urban-E | Urban-W |
|---|---|---|---|---|---|---|---|---|---|
| N % | Site | 3 | 5.779 | 0.0011* | mean | 0.62% | 0.82% | 0.48% | 0.76% |
| log10 | Depth | 1 | 0.2103 | 0.6475 | SE | 0.01 | 0.02 | 0.00 | 0.01 |
| | Site x Depth | 3 | 1.06 | 0.3698 | groups | ab | b | a | b |
| C:N | Site | 3 | 3.3989 | 0.0207* | mean | 28.24 | 24.77 | 31.56 | 25.02 |
| | Depth | 1 | 0.0687 | 0.7938 | SE | 0.4 | 0.4 | 0.23 | 0.31 |
| | Site x Depth | 3 | 1.2926 | 0.2811 | groups | ab | a | b | a |
| Chlorodesmis sp. | | | | | | | | | |
| N % | Site | 1 | 6.0924 | 0.0261* | mean | 2.63% | - | 3.31% | 2.36% |
| | Depth | 1 | 0.3489 | 0.5635 | SE | 0.1 | | 0.05 | 0.1 |
| | Site x Depth | 1 | 0.3849 | 0.5443 | groups | | | b | a |
| C:N | Site | 1 | 4.2869 | 0.0561 | mean | 11.78 | - | 9.71 | 10.95 |
| | Depth | 1 | 6.0953 | 0.0261* | SE | 0.5 | | 0.06 | 0.23 |
| | Site x Depth | 1 | 1.6411 | 0.2196 | groups | | | a | a |

Table A7 PERMANOVA analyses testing the effect of Site, Depth, and seawater nutrient concentrations ($NH_4^+$, $NO_3^-$, and $PO_4^{3-}$) on the of the prevalence of coral disease and signs of compromised coral health from surveys in Timor-Leste in 2015. Bolded values are significant results with significance at p = 0.05. df–degrees of freedom

| **Source of Variation** | **df** | **Sums of Squares** | **Means of Squares** | **Pseudo-F** | **$r^2$** | **p(perm)** |
|---|---|---|---|---|---|---|
| Site | 3 | 0.52615 | 0.175383 | 7.4077 | 0.42076 | 0.0001* |
| Depth | 1 | 0.12196 | 0.121964 | 5.1514 | 0.09753 | 0.0016* |
| $NH_4$+ | 1 | 0.04585 | 0.045849 | 1.9365 | 0.03667 | 0.1002 |
| $NO_3$- + $NO_2$- | 1 | 0.01789 | 0.017892 | 0.7557 | 0.01431 | 0.5945 |
| $PO_4^{3-}$ | 1 | 0.07698 | 0.076982 | 3.2515 | 0.06156 | 0.0162* |
| Site x Depth | 3 | 0.15385 | 0.051284 | 2.1661 | 0.12303 | 0.0220* |

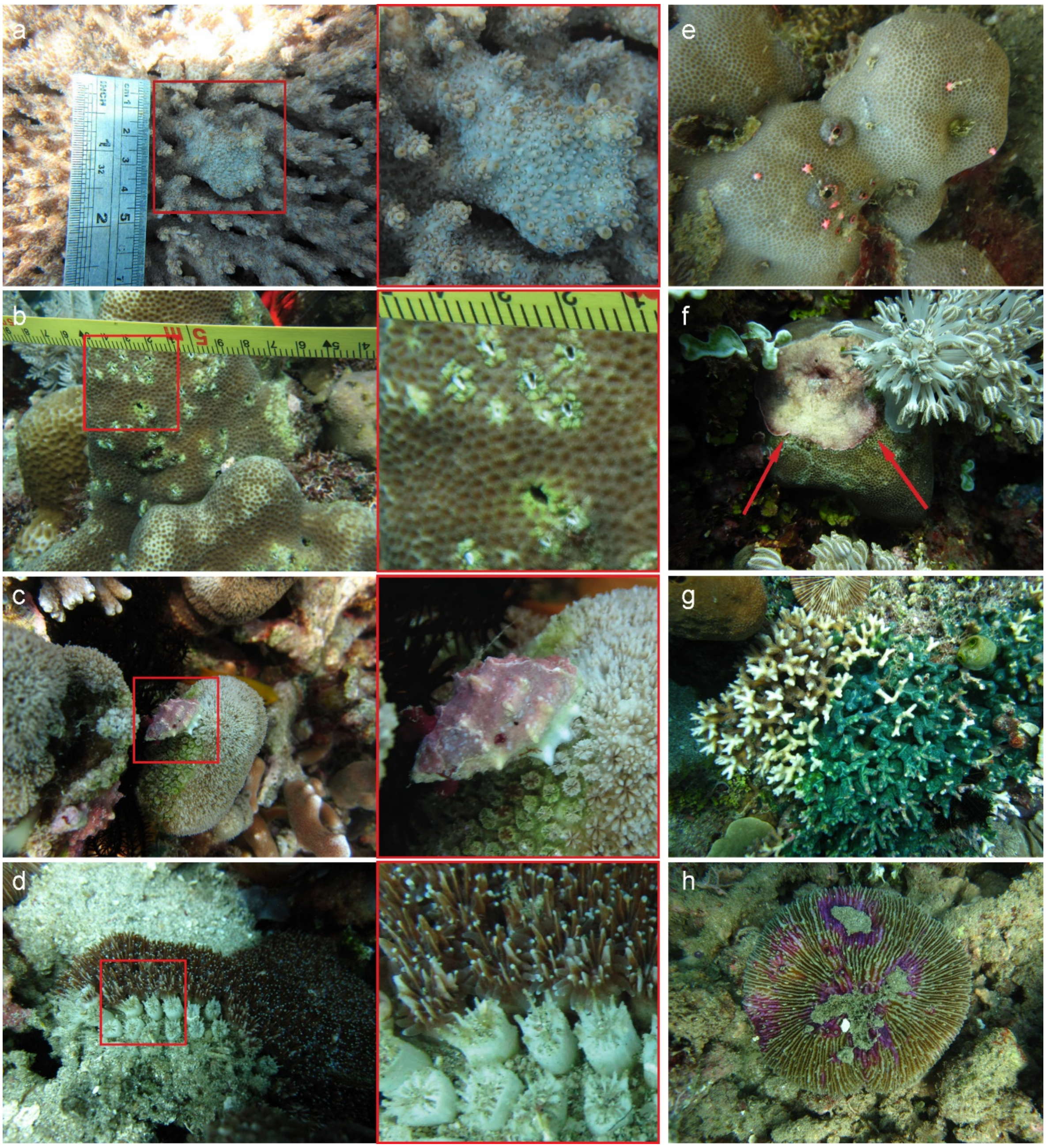


Figure A1 Other diseases and compromised states found during surveys: a) GA-growth anomalies, b) burrowing barnacles, c) *Drupella sp.* snail predation, d) TL-unexplained tissue loss, e) potential Trematodiasis, f) CCA overgrowth, g) tunicate overgrowth, and h) pigmentation. See Table A3 for more details.

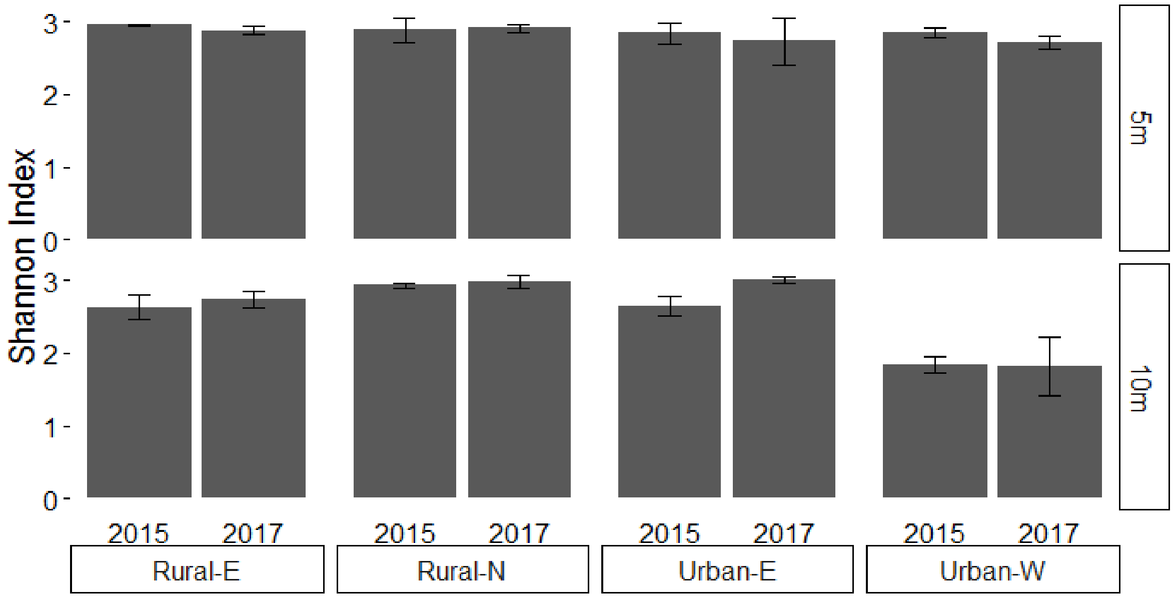


Figure A2 Shannon Diversity Index calculated per site and depth on genera present per transect in Timor-Leste. Top bars are at 5 m depth with 10 m below. Sites on the x-axis are split into the 2015 and 2017 survey periods. Error bars show standard error.

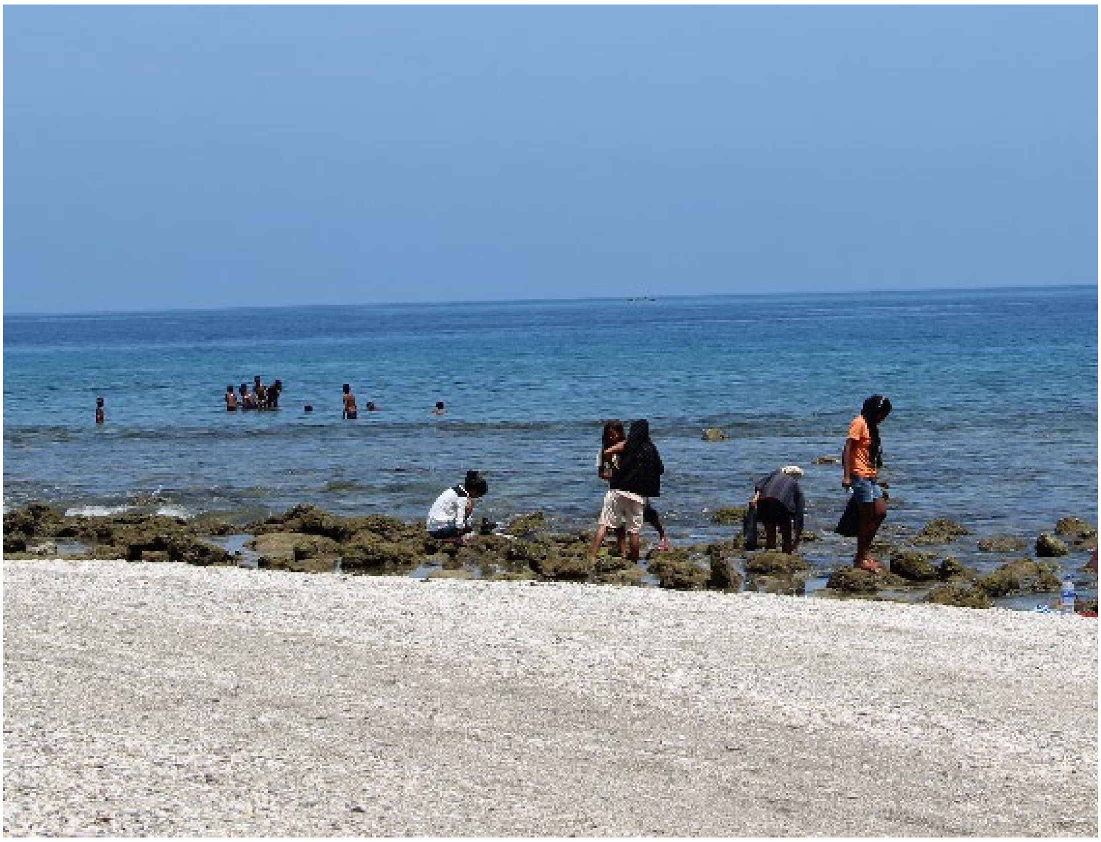

Figure A3 Gleaning and recreational activity at Urban-W site in Timor-Leste during 2015 surveys. Urban-W is in the Dom Alexio subdistrict of Dili which had a population density of 5017.9 people per km$^2$ in the 2015 Timor-Leste census. Comparatively, the national population density is 79.3 people per km$^2$. This beach was within walking distance to densely populated neighborhoods and very popular for swimming, fishing, and gleaning which likely has impacted the shallow-water coral reefs. Women can be seen in the foreground gleaning and children playing on the reef flat further offshore.

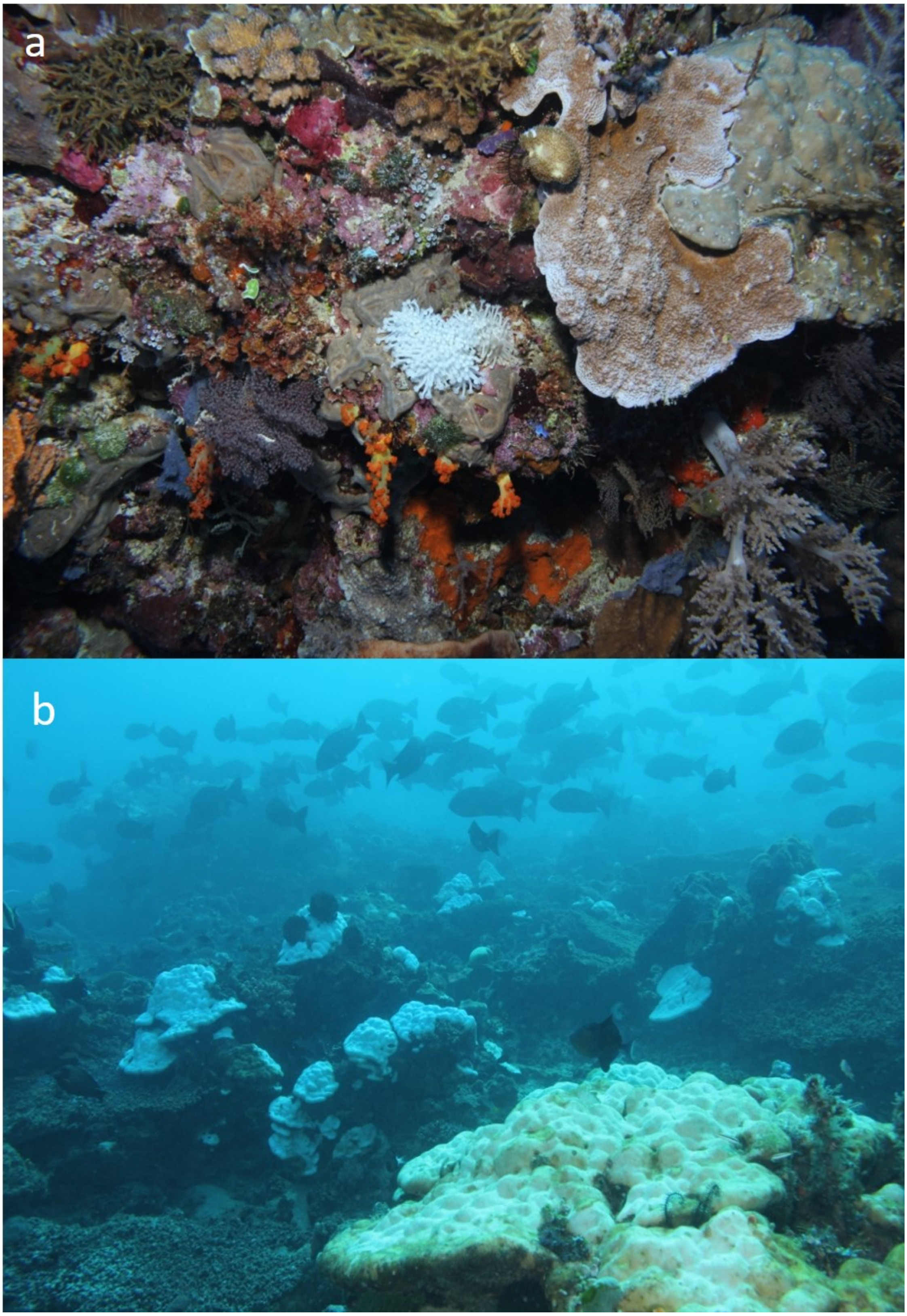


Figure A4 Photos by Tony Crean, a local dive operator, emailed on May 31, 2016. a) Bleached *Goniopora sp.* at Adara on the west coast of Ataúro Island. An estimated 90% of *Goniopora sp.* were bleached with no other hard corals affected on Ataúro Island reefs. Multiple other genera of corals are visibly unbleached. Depth of photo unknown. b) Image was taken at Jaco Island, the easternmost point of Timor-Leste with bleaching of massive *Porites sp.* corals. Similar percent of corals bleached from 5–18 m.

**Appendix:**

Nutrients measured in the Laclo river in 2006 indicated low levels of pollution from sewage and nutrients from fertilizer [20]. The report concluded that nutrient levels in Laclo were comparable to other tropical rivers [197–199] and tidal seawater samples were similar to unpolluted mangrove sampling in Australia [200]. Ammonium decreased from 4 μM approximately 10 km upstream from the mouth of the Laclo, to ~2 μM at the mouth, and then to <1 μM about 20 km west of the mouth in Metinaro. Combined nitrate and nitrite dropped from >8 μM to almost zero from the river sampling to Metinaro. There was a similar pattern with phosphate concentrations of 0.5 μM in the Laclo which dropped to 0.05 μM in Metinaro (Table A8). In the present study, the site Rural-E at Manatuto was approximately half-way (i.e., 12 km from the mouth of the Laclo) between the mouth of the Laclo River and Metinaro. Seawater nutrient samples at Rural-N 5 m were in-between values sampled at the Laclo River and Metinaro (Table A8). As there are 10 years, riverine and coastal versus sampling at 5 m depth on the reef between the Laclo and Rural-E samplings and it is difficult to draw conclusions based on these limited nutrient measurements. Additionally, Rural-E is about 12 km west of the mouth of the Laclo and how much riverine outputs affect the downstream coast is not known. However, both surveys were taken during the dry season and based on the limited evidence it does not appear that nutrients on the reef at Rural-E are greatly elevated by anthropogenic inputs.

Table A8 Nutrients from water samples collected in the Laclo River and Metinaro in 2006 as reported in [20] and from Rural-E at 5 m depth in 2015 which is approximately half-way between the mouth of the Laclo and Metinaro in Timor-Leste.

| Site | $NH_4^+$ μM | $NO_2^- + NO_3^-$ μM | $PO_4^{3-}$ μM |
|---|---|---|---|
| Laclo River Upstream | 4.04 ± 0.93 | 8.22 ± 0.38 | 0.49 ± 0.09 |
| Laclo River Mouth | 1.98 ± 1.03 | 8.65 ± 1.32 | 0.54 ± 0.01 |
| Rural-E at 5 m (12 km from mouth) | 1.20 ± 0.28 | 0.90 ± 0.16 | 0.11 ± 0.01 |
| Metinaro (20 km from mouth) | 0.80 ± 0.09 | 0.05 ± 0.00 | 0.05 ± 0.01 |
